# Habitual control of instrumental behavior driven by corticostriatal pathways originating from the primary motor cortex

**DOI:** 10.64898/2026.11.25.769086

**Authors:** Shigeki Kato, Nozomu Yoshioka, Yoshio Iguchi, Masateru Sugawara, Kayo Nishizawa, Susumu Setogawa, Yuko Maejima, Kenju Shimomura, Masahiro Okamoto, Satoshi Eifuku, Kazuto Kobayashi

**Author notes:** Nozomu Yoshioka, Division of Functional Morphology, Department of Anatomy, Iwate Medical University, Iwate, Japan. **Competing Interest Statement:** The authors have no competing interests to declare.

## Abstract

Instrumental performance is governed by goal-directed and habitual modes of behavioral control, with their relative contributions shifting during habit formation. These two modes of control involve distinct regions of the dorsal striatum: the dorsomedial striatum (DMS) mediates goal-directed control, whereas the dorsolateral striatum (DLS) drives habitual control. Although corticostriatal inputs from the frontal cortices to the DMS have been shown to support goal-directed control, which cortical inputs to the DLS underlie habitual control remains unclear. Here, we selectively eliminated corticostriatal neurons originating from either the primary motor cortex (M1) or primary somatosensory cortex (S1) and projecting to the DLS in mice. The balance between goal-directed and habitual control in a food-reinforced lever-press task was evaluated using outcome devaluation and contingency degradation. Selective elimination of corticostriatal neurons originating in M1, but not S1, compromised the insensitivity of lever pressing to outcome devaluation and contingency degradation, suggesting impaired habit formation. Chemogenetic silencing of the M1–DLS corticostriatal neurons also impaired habit formation. Furthermore, we investigated the functional roles of pyramidal tract (PT) and intratelencephalic (IT) neuronal populations constituting the M1–DLS pathway. Selective elimination of PT neurons impaired insensitivity to outcome devaluation without significantly affecting performance following contingency degradation, whereas selective elimination of IT neurons had no detectable effect in either test. Our findings suggest that the M1–DLS pathway plays an important role in habitual control and that PT neurons contribute, at least in part, to the insensitivity to outcome devaluation associated with habit formation.

**Significance Statement:** The dorsolateral striatum (DLS) mediates habitual control, which can optimize behavioral execution and reduce cognitive demands. However, the precise functional contributions of distinct corticostriatal inputs to habitual control remain unclear. We demonstrated that the corticostriatal pathway from the primary motor cortex to the DLS (M1–DLS pathway), but not that from the primary somatosensory cortex, is essential for the development of habitual behavior in mice. We further found that pyramidal tract neurons, but not intratelencephalic neurons, within the M1–DLS pathway are required for the insensitivity to outcome devaluation that accompanies habit formation, indicating functional diversity among projection-defined neuronal populations. These findings suggest that the corticostriatal system may represent a therapeutic target for neuropsychiatric disorders characterized by impaired habit formation.

## Introduction

Instrumental learning enables animals to act on their environment to control the occurrence of biologically significant events. Instrumental performance is governed by goal-directed and habitual modes of behavioral control (1, 2). During the early phase of conditioning, it is primarily goal-directed, based on action-outcome associations, allowing flexible adaptation but requiring greater cognitive effort (3–5). With extended training, however, performance can come under habitual control based on stimulus-response associations, enabling efficient behavioral execution at the expense of flexibility (5–7). The growing dominance of habitual over goal-directed control is considered to underlie habit formation (8, 9).

The cortico-basal ganglia loops involving the dorsal striatum play a key role in instrumental learning and decision-making (10–12). The dorsal striatum can be subdivided into the dorsomedial striatum (DMS) and dorsolateral striatum (DLS), which receive topographically distinct afferent projections from different cortical areas. The DMS mainly receives projections from the prelimbic cortex (PrL), orbitofrontal cortex (OFC), and secondary motor cortex (M2), whereas the DLS receives predominant inputs from the primary motor cortex (M1) and primary somatosensory cortex (S1) (13, 14). Previous studies with excitotoxic lesions and pharmacological inactivation of the DMS or DLS have demonstrated that these regions contribute to goal-directed and habitual control of instrumental behavior, respectively (8, 15, 16). Subsequent studies have reported that corticostriatal inputs from the PrL or OFC to the DMS mediate the acquisition and context-dependent expression of goal-directed actions (17–19). The firing properties of DLS neurons in response to corticostriatal inputs are correlated with the degree of habitual behavior (20). However, the specific cortical inputs to the DLS that drive habit formation remain unclear.

Corticostriatal neurons are located mainly in layers II/III and V of the cerebral cortex and are classified primarily into pyramidal tract (PT) neurons and intratelencephalic (IT) neurons, distinguished by their projection patterns, electrophysiological properties, and gene expression profiles (21, 22). PT neurons send axonal and collateral projections to the ipsilateral striatum, subthalamic nucleus, thalamus, superior colliculus, and brainstem (23, 24), whereas IT neurons project to striatal and cortical areas in both hemispheres but not to regions outside the telencephalon (25, 26). Recent evidence suggests functional differences between PT and IT neurons in movement, drug response, and instrumental learning (27–29). However, whether PT and IT neurons differentially support habit formation remains largely unexplored.

In the present study, we investigated the roles of corticostriatal neurons originating from M1 or S1 and projecting to the DLS in habit formation during instrumental learning in mice. Selective targeting of these pathways indicated that M1, but not S1, corticostriatal neurons underlie habitual responses. Chemogenetic silencing of the M1-derived corticostriatal pathway provided further evidence of its role in habitual control. Furthermore, we studied the involvement of PT and IT neurons in the M1–DLS pathway to habitual control. Selective manipulation of these neuronal populations suggested that PT neurons contribute, at least in part, to habit formation.

## Results

### Projection patterns of M1- and S1-derived corticostriatal neurons

The dorsal striatum receives dense innervation from various cerebral cortical areas (13, 14). To characterize the projection patterns of corticostriatal neurons originating from M1 and S1 and projecting to the DLS, we employed a double-vector transduction strategy (Fig. 1A). In this strategy, a neuron-specific retrograde gene transfer (NeuRet) vector (30) carrying a Cre transgene was unilaterally injected into the dorsal striatum to deliver the transgene to corticostriatal neurons. Subsequently, adeno-associated viral (AAV) vectors carrying red fluorescent protein (RFP) and green fluorescent protein (GFP) transgenes under the control of the flip-excision switch (FLEX) system (31–33) were injected into M1 and S1, respectively, ipsilateral to the NeuRet-Cre vector injection to achieve differential labeling of these corticostriatal neurons (see *SI Appendix*, Fig. S1 *A* and *B* for the coordinates of vector injections). Retrograde delivery of the Cre transgene to multiple cortical areas was confirmed by immunohistochemistry with an anti-Cre antibody (*SI Appendix*, Fig. S2). FLEX-mediated RFP and GFP fluorescence signals were visualized in both hemispheres (Fig. 1B) and were observed in the cell bodies of corticostriatal neurons in the ipsilateral M1 and S1, respectively (Fig. 1C). RFP- and GFP-positive signals were detected in nerve terminals innervating the DLS in both hemispheres (Fig. 1D), as well as in terminals projecting to contralateral cortical areas (Fig. 1E). Axonal collaterals of these corticostriatal neurons were also labeled in the contralateral superior colliculus and in brainstem regions, including the pontine nucleus (Pn) and the pyramidal tract of the medulla oblongata (*SI Appendix*, Fig. S3).

**Fig. 1.**
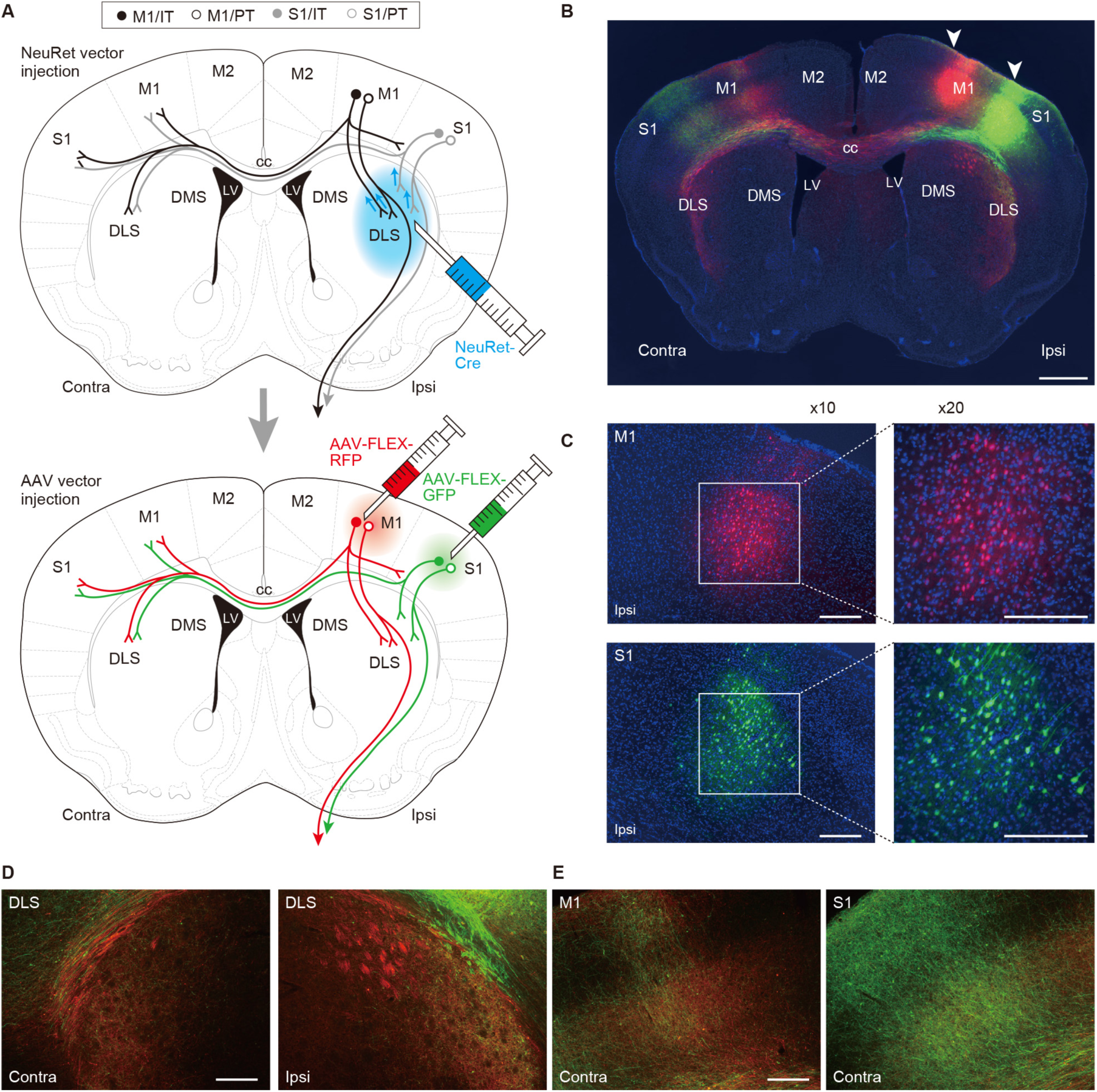
Distribution of the M1–DLS and S1–DLS corticostriatal pathways. (*A*) Experimental strategy for pathway labeling. The NeuRet vector encoding Cre recombinase was unilaterally injected into the mouse DLS, from which it was retrogradely transported to the cell bodies projecting to the DLS. AAV-FLEX-RFP and AAV-FLEX-GFP vectors were then injected into M1 and S1, respectively, ipsilateral to the NeuRet vector injection. Consequently, RFP and GFP were selectively expressed in corticostriatal neurons innervating the DLS. (*B*) A representative coronal section showing fluorescently labeled corticostriatal pathways. (*C*) Magnified views of the injection-side M1 and S1, showing fluorescently labeled pyramidal neurons. (*D*) Detection of RFP- and GFP-positive axon terminals in the DLS contralateral and ipsilateral to the injections. (*E*) Labeled axon terminals in the contralateral M1 and S1. Contra, contralateral; Ipsi, ipsilateral. (Scale bars: *B*, 1 mm; *C*–*E*, 200 μm.)

### Selective elimination of M1- and S1-derived corticostriatal neurons

To address the behavioral roles of corticostriatal neurons innervating the DLS, we used a selective neural pathway targeting technology (34, 35) to eliminate those originating from either M1 or S1 in the mouse brain. The NeuRet vector carrying a transgene encoding the human interleukin-2 receptor α-subunit fused to enhanced GFP (IL-2Rα-GFP) was bilaterally injected into the DLS. Four weeks later, either a solution containing a recombinant immunotoxin (ITX) (36, 37) or 0.1 M phosphate-buffered saline (PBS; pH 7.4) was unilaterally injected into M1 or S1 (see *SI Appendix*, Fig. S1*C* for the coordinates of ITX or PBS injections). Brain sections through M1 and S1 were subjected to GFP immunohistochemistry (see *SI Appendix*, Fig. S4 for sections used for cell counts). Following ITX or PBS injection into M1, fewer GFP-positive corticostriatal neurons were observed in the M1 on the ITX-injected side than on the PBS-injected side, whereas no corresponding reduction was observed in the S1 (Fig. 2A). Quantification showed a significantly lower number of GFP-positive neurons in the ITX-injected M1 than in the PBS-injected M1 (Fig. 2B). Similarly, following ITX or PBS injection into the S1, fewer GFP-positive neurons were observed in the S1 on the ITX-injected side than on the control side (Fig. 2D), and cell counts confirmed a significant reduction in the S1 (Fig. 2E). Cresyl violet staining revealed no apparent tissue damage in the ITX-injected cortical areas (Fig. 2 *C* and *F*). These results indicate that ITX injection selectively eliminated retrogradely labeled corticostriatal neurons originating in M1 or S1.

**Fig. 2.**
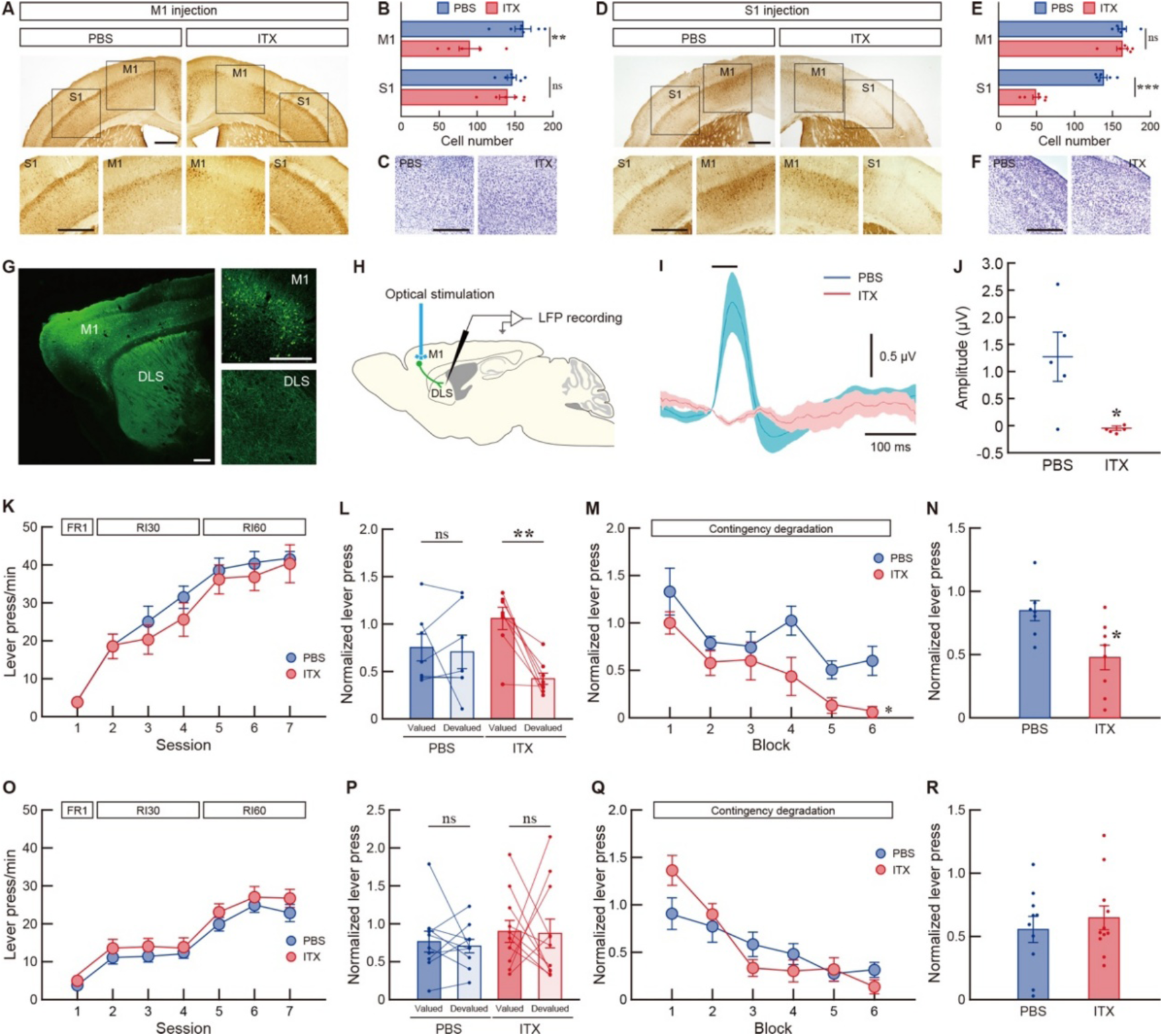
Selective elimination of corticostriatal neurons and its effects on habit formation. (*A*–*F*) Histological validation of cell elimination. Mice received bilateral injections of the NeuRet-IL-2Rα-GFP vector into the DLS, followed by a unilateral injection of ITX into M1 or S1, with PBS injected into the contralateral hemisphere as a control. (*A*) Coronal sections containing the M1 and S1 from mice following PBS or ITX injection into the M1, stained with an anti-GFP antibody. The number of GFP-positive cells in M1 was lower on the ITX-injected side than on the PBS-injected side, whereas that in the S1 was similar between the two sides. (*B*) Numbers of GFP-positive cells per section in the M1 and S1 (*n* = 6 samples per group). (*C*) Cresyl violet staining of M1 sections on the PBS- and ITX-injected sides. (*D*) Sections containing the M1 and S1 from mice following PBS or ITX injection into the S1, stained with an anti-GFP antibody. The number of GFP-positive cells in the S1 was lower on the ITX-injected side than on the PBS-injected side, whereas that in M1 was comparable between the two sides. (E) Numbers of GFP-positive cells per section in the M1 and S1 (*n* = 6 per group). (F) Cresyl violet staining of S1 sections on the PBS- and ITX-injected sides. (*G*– *J*) Electrophysiological recordings of neuronal activity evoked by optical stimulation. For neuronal recordings, mice received an injection of the NeuRet-Cre vector into the DLS, followed by an injection of the AAV-FLEX-IL-2Rα-2A-ChR2-YFP vector into the M1 and a subsequent injection of PBS or ITX into M1. (G) Expression of IL-2Rα-2A-ChR2-YFP in corticostriatal neurons of the PBS-injected mice. (*H*) Experimental diagram for recording neuronal activity evoked by optical stimulation. Blue light was delivered to M1, and LFPs were recorded in the DLS. (*I*) LFPs in the DLS evoked by optogenetic stimulation of the M1–DLS pathway. The black bar indicates a 50-ms blue light pulse. Grand-averaged waveforms from PBS- and ITX-injected mice are shown, with negative voltages plotted upward. The peak amplitude was significantly lower in the ITX-treated group than in the PBS-injected group. (*J*) Peak amplitudes of LFP responses recorded from the DLS (*n* = 5 per group). (*K*–*R*) Behavioral assessment of habit formation. Mice received bilateral injections of the NeuRet-IL-2Rα-GFP vector into the DLS, followed by injections of PBS or ITX into M1 or S1, and were subjected to a series of lever-press tasks. (*K*) Acquisition of the lever-press task in mice with M1 injections. (*L*) Outcome devaluation test in mice with M1 injections. The number of lever presses during a 5-min test session under valued and devalued states was normalized to the number of lever presses in session 7. (*M*) Contingency degradation test in mice with M1 injections. The number of lever presses in each 5-min block was normalized to the number of lever presses session 12. (*N*) Session mean of the normalized number of lever presses in mice with M1 injections. (*O*) Acquisition of the lever-press task in mice with S1 injections. (*P*) Outcome devaluation test for mice with S1 injections. (*Q*) Contingency degradation test in mice with S1 injections. (R) Session mean of the normalized number of lever presses in mice with S1 injections. For K–R, M1 injections: *n* = 7 mice (PBS), *n* = 8 mice (ITX); S1 injections: *n* = 10 mice (PBS), *n* = 11 mice (ITX). Data in *B*, *E, J,* and *K*–*R* are presented as mean ± SEM; Statistical analyses were performed using unpaired *t*-test (*B*, *E, J, N* and *R*), two-way ANOVA followed by pairwise comparisons with Bonferroni correction (*K*, *M, O* and *Q*), or planned comparisons using paired *t*-test (*L* and *P*). See also *SI Appendix*, Table S1 for statistical details. \**P* < 0.05; \*\**P* < 0.01; \*\*\**P* < 0.001; ns, not significant. (Scale bars: *A*, *C*, *D*, *F*, *and G,* 200 μm.)

To assess whether the selective elimination of corticostriatal neurons alters functional connectivity between M1 and the DLS, we recorded neural responses in the DLS evoked by optical stimulation of M1. The NeuRet-Cre vector was unilaterally injected into the dorsal striatum, and an AAV vector carrying a gene cassette encoding IL-2Rα, a 2A peptide, and channelrhodopsin-2 (H134R) fused to yellow fluorescent protein (ChR2-YFP) under the control of the FLEX system was injected into the ipsilateral M1. Subsequently, ITX or PBS was injected into M1 ipsilateral to the vector injections. GFP immunostaining detected transgene expression in the cell bodies of M1 corticostriatal neurons and their DLS nerve terminals in PBS-injected mice (Fig. 2G). Flashes of 465-nm light were delivered to M1 through an optical fiber, while local field potentials (LFPs) were recorded in the DLS (Fig. 2H). Following the recording sessions, the placement of the optical fibers and recording electrodes was histologically verified (*SI Appendix*, Fig. S5). In the PBS-injected mice, LFP responses were evoked approximately 15 ms after optical stimulation; in contrast, this LFP component was nearly abolished in the ITX-injected mice (Fig. 2I). The average peak amplitude of the LFP responses in the ITX-injected mice was significantly smaller than that in the PBS-injected mice (Fig. 2J). Taken together, these electrophysiological findings demonstrate that selective elimination of M1-derived corticostriatal neurons markedly reduced M1-evoked LFP responses in the DLS.

### Elimination of corticostriatal neurons in the M1–DLS pathway impaired habit formation

We investigated the effects of selective elimination of corticostriatal neurons innervating the DLS on habit formation. The relative contributions of goal-directed and habitual control were assessed during a food-reinforced lever-press task using outcome devaluation and contingency degradation paradigms (38–40). Performance under goal-directed control is sensitive to changes in the current incentive value of the outcome and the action-outcome contingency, whereas performance under habitual control is relatively insensitive to these changes. Mice received bilateral intracranial injections of the NeuRet-IL-2Rα-GFP vector into the DLS, followed by unilateral injections of ITX or PBS into either M1 or S1. They then underwent a series of operant lever-press tasks reinforced with 20-mg food pellets. Mice were trained on a fixed ratio 1 (FR1) schedule on day 1 (session 1), followed by random-interval (RI) schedules with average intervals of 30 s (RI30) and 60 s (RI60). Each RI schedule was maintained for 3 days (sessions 2–4 for RI30 and sessions 5–7 for RI60).

For the outcome devaluation test, mice were given *ad libitum* access to either standard laboratory chow (valued state) or the food pellets used as the reinforcer (devalued state) for 1 h, followed immediately by a 5-min lever-press test conducted without reinforcement (session 8). On the next day, the mice underwent the opposite prefeeding condition (valued or devalued) and tested under the same 5-min nonreinforced conditions (session 9). The number of lever presses during these test periods was normalized to the number of lever presses recorded in session 7. After the devaluation test, mice underwent 3 days of retraining under the RI60 schedule (sessions 10–12). During the 30-min contingency degradation test, which was divided into six 5-min blocks, a food pellet was delivered only when 20 s had passed without any lever presses. Each lever press reset the timer, thereby delaying pellet delivery. The number of lever presses in each test block was normalized to the number of lever presses recorded in session 12.

Following PBS or ITX injection into the M1, both the PBS- and ITX-injected groups exhibited a gradual and comparable increase in the number of lever presses across conditioning sessions 1–7 (Fig. 2K). In the outcome devaluation test, the normalized lever press rate in the PBS-injected group was comparable between the valued and devalued states, indicating insensitivity to outcome devaluation. In contrast, the ITX-injected group showed a significant reduction in lever presses in the devalued state compared with the valued state, indicating a disruption of insensitivity to outcome devaluation (Fig. 2L). In the contingency degradation test, normalized lever presses were lower across blocks in the ITX-injected group than in the PBS-injected group (Fig. 2M). Furthermore, the overall average number of normalized lever presses was significantly lower in the ITX-injected group than in the PBS-injected group (Fig. 2N), indicating greater sensitivity to contingency degradation in the ITX-injected group. Collectively, these results suggest that selective targeting of M1-derived corticostriatal neurons impairs habit formation.

Following PBS or ITX injection into the S1, both the PBS- and ITX-injected groups showed a similar increase in lever presses during the conditioning sessions (Fig. 2O). In the outcome devaluation test, the normalized number of lever presses in both groups did not differ significantly between the valued and devalued states (Fig. 2P). In the contingency degradation test, the normalized number of lever presses showed a similar pattern across blocks in the PBS- and ITX-injected groups (Fig. 2Q), with no significant difference in the session averages between the groups (Fig. 2R). Unlike selective elimination of M1-derived corticostriatal neurons, selective elimination of S1-derived corticostriatal neurons did not impair habit formation.

Because the DLS is implicated in spontaneous locomotion and motor coordination (41, 42), we also assessed whether the elimination of M1- or S1-derived corticostriatal neurons affected basic motor performance. Spontaneous locomotor activity was monitored in an open field for 60 min, and motor coordination was assessed by measuring the latency to fall from an accelerating rotarod. Following M1 injections, spontaneous locomotor activity showed no significant difference between the PBS- and ITX-injected groups (*SI Appendix*, Fig. S6*A*). Furthermore, the latency to fall increased progressively in both groups, with no significant difference between the groups (*SI Appendix*, Fig. S6*B*). Similarly, following S1 injections, no significant differences were found between the groups in either spontaneous locomotor activity (*SI Appendix*, Fig. S6*C*) or the latency to fall (*SI Appendix*, Fig. S6*D*). These results indicate that selective elimination of M1- or S1-derived corticostriatal neurons did not significantly affect spontaneous locomotion, motor coordination, or motor skill learning, as assessed by these tests. Thus, the impairment of habit formation caused by selective elimination of M1-derived corticostriatal neurons is unlikely to be explained by deficits in basic motor performance.

### Chemogenetic silencing of the M1–DLS pathway impaired habit formation

To further assess the behavioral roles of corticostriatal neurons in the M1–DLS pathway, we performed chemogenetic silencing of this pathway using a modified invertebrate glutamate-gated chloride channel composed of α- and β-subunits (GluClαβ). In this variant, a mutation in the β-subunit abolishes responsiveness to endogenous glutamate while maintaining high sensitivity to the anthelmintic drug ivermectin (IVM) (43). To express GluClαβ specifically in corticostriatal neurons, mice received bilateral injections of the NeuRet-Cre vector into the DLS, followed by injections of AAV vectors into M1 encoding GluClα fused to mYFP (GluClα-mYFP) and GluClβ fused to RFP (GluClβ-RFP) under the control of the FLEX system. Double immunohistochemistry with anti-GFP and anti-RFP antibodies detected the coexpression of both subunits in numerous M1 neurons (Fig. 3A). To validate the functional efficacy of this system, we performed whole-cell patch-clamp recordings in acute brain slices containing M1 from vector-injected mice. Bath application of IVM (20 µM) suppressed action potential firing in GluClαβ-expressing neurons (Fig. 3B), with significant membrane hyperpolarization and a significant reduction in firing frequency (Fig. 3C).

**Fig. 3.**
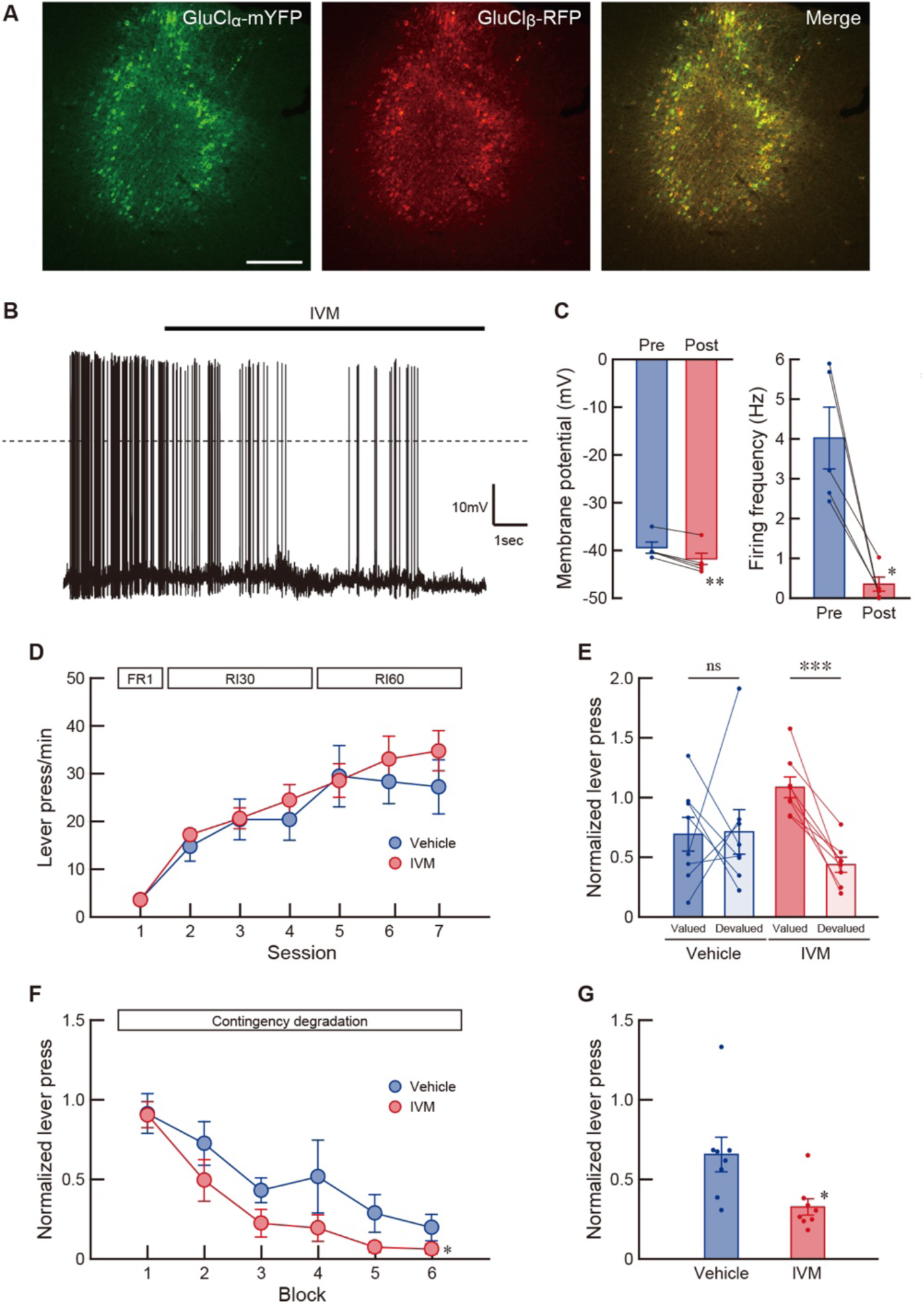
Chemogenetic silencing of M1–DLS corticostriatal neurons and its effects on habit formation. Mice received bilateral injections of the NeuRet-Cre vector into the DLS, followed by injections of AAV-FLEX-GluClα-mYFP and AAV-FLEX-GluClβ-RFP vectors into M1. (*A*) Histological validation. Expression of GluClαβ in M1-DLS corticostriatal neurons. Sections through M1 were subjected to double immunohistochemistry with anti-GFP and anti-RFP antibodies. (*B* and *C*) Electrophysiological verification of chemogenetic silencing. (*B*) Whole-cell patch-clamp recordings from slice preparations. IVM (20 µM) was applied to assess its effect on neuronal firing activity. A representative firing pattern of a neuron is shown. The dashed line represents 0 mV. (*C*) Membrane potentials and firing frequencies of GluClαβ-expressing neurons (*n* = 5 per group). (*D*–*G*) Behavioral assessment in mice receiving intraperitoneal injection of IVM (5.0 mg/kg) or vehicle. (*D*) Acquisition of the lever-press task. (*E*) Outcome devaluation test. The number of lever presses during a 5-min test session under valued and devalued states was normalized to the number of lever presses in session 7. (*F*) Contingency degradation test. The number of lever presses during a 5-min test block was normalized to the number of lever presses in session 12. (*G*) Session mean of the normalized number of lever presses (*n* = 8 per group). Data in *C–G* are presented as mean ± SEM; Statistical analyses were performed using paired *t*-test (*C* and *E*), two-way repeated-measures ANOVA (*D* and *F*), or an unpaired *t*-test (*G*). See also *SI Appendix*, Table S1 for statistical details. \**P* < 0.05; \*\**P* < 0.01; \*\*\**P* < 0.001; ns, not significant. (Scale bar: *A,* 200 μm.)

For behavioral assessment, vector-injected mice were subjected to the same series of lever-press tasks as those used for mice subjected to selective elimination of M1–DLS corticostriatal neurons. During the behavioral testing period, mice received intraperitoneal injections of either IVM (5.0 mg/kg) or vehicle immediately after the FR1 session and every other day thereafter. In both the vehicle- and IVM-injected mice, the number of lever presses gradually increased as the sessions progressed, with similar acquisition patterns between the groups (Fig. 3D). In the outcome devaluation test, the normalized number of lever presses in the vehicle-injected group did not differ significantly between the valued and devalued states, indicating insensitivity to outcome devaluation, whereas the IVM-injected group displayed a significant reduction in lever pressing in the devalued state compared with the valued state, indicating an impairment of insensitivity to outcome devaluation (Fig. 3E). In the contingency degradation test, the normalized number of lever presses was lower across blocks in the IVM-injected group than in the vehicle-injected group (Fig. 3F). The session average was significantly lower in the IVM-injected group than in the vehicle-injected group (Fig. 3G), indicating greater sensitivity to contingency degradation in the IVM-injected group than in the vehicle-injected group. Taken together, these findings indicate that chemogenetic silencing of the M1–DLS corticostriatal pathway impaired habit formation, consistent with the effects of selective elimination of M1-derived corticostriatal neurons.

### Differential influences of selective elimination of PT and IT corticostriatal neurons on habit formation

To determine which neuronal type within the M1–DLS corticostriatal pathway— PT or IT—is involved in habit formation, we selectively eliminated each neuronal population. To selectively eliminate PT neurons, the NeuRet-IL-2Rα-GFP and NeuRet-RFP vectors were unilaterally injected into the Pn and the contralateral DLS, respectively. Subsequently, ITX or PBS was injected into the M1 ipsilateral to the NeuRet-IL-2Rα-GFP injection (Fig. 4A). Immunohistochemical staining of M1 sections showed a marked reduction in the number of GFP-positive PT neurons in the ITX-injected group compared with that in the PBS-injected group, whereas no apparent difference was observed in the number of RFP-positive IT neurons between the two groups (Fig. 4B). Consistent with this observation, the number of PT neurons was significantly lower in the ITX-injected group than in the PBS-injected group (Fig. 4C).

**Fig. 4.**
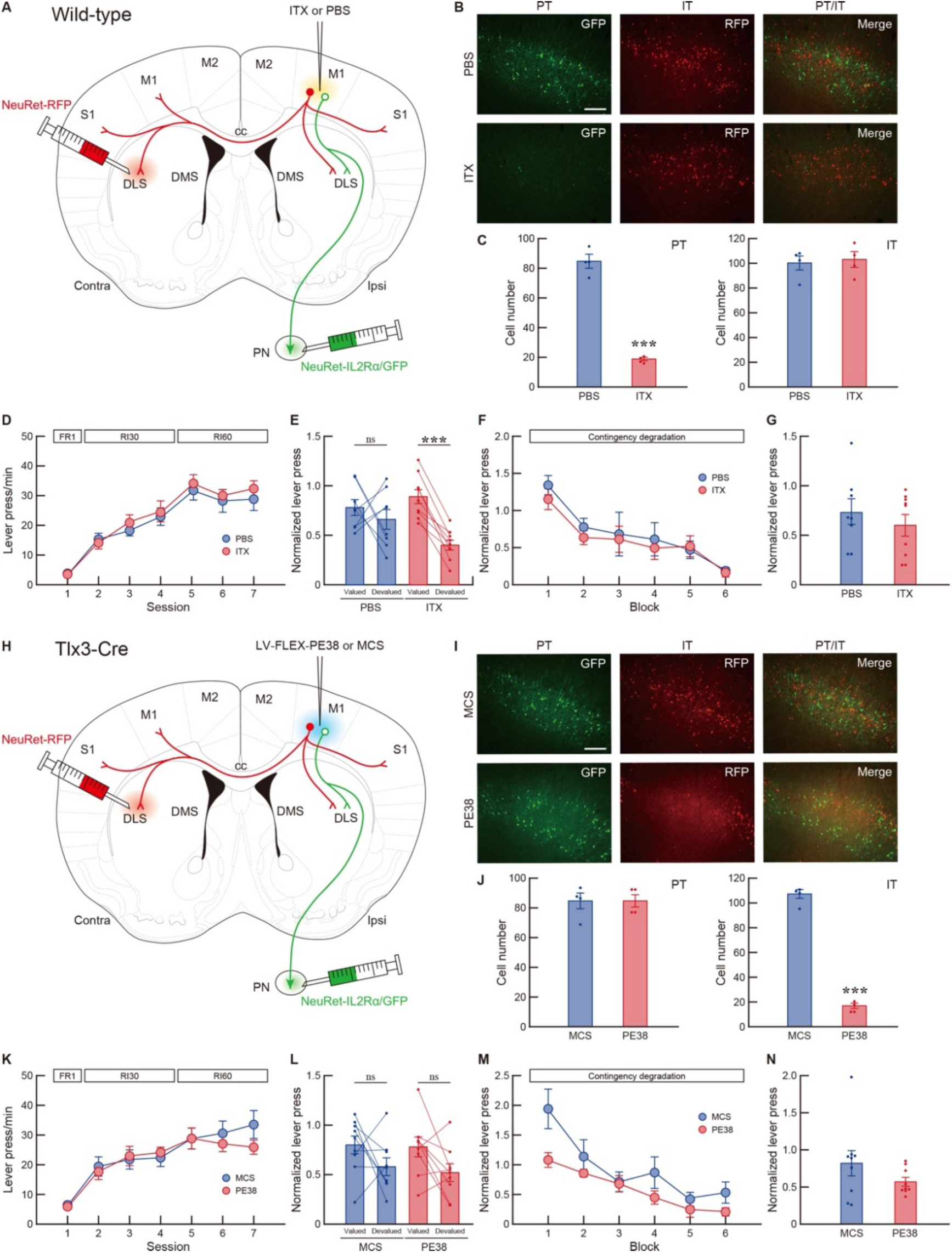
Selective elimination of PT and IT corticostriatal neurons and its effects on habit formation. (*A*–*G*) Selective elimination of M1–DLS PT neurons and assessment of its behavioral effects. (*A*) Experimental strategy. NeuRet-IL-2Rα-GFP and NeuRet-RFP vectors were unilaterally injected into the Pn and the contralateral DLS, respectively, followed by an injection of PBS or ITX into M1 ipsilateral to the NeuRet-IL-2Rα-GFP injection. (*B*) Distribution of PT and IT neurons in M1. Sections through M1 were subjected to double immunohistochemistry with anti-GFP and anti-RFP antibodies. (*C*) Numbers of PT and IT neurons per section (*n* = 4 per group). (*D*–*G*) Behavioral assessment of C57BL/6J mice following M1–DLS PT neuron elimination. Mice received bilateral injections of the NeuRet-IL-2Rα-GFP vector into the Pn, followed by injections of PBS or ITX injections into M1 and subjected to a series of lever-press tasks. (*D*) Acquisition of the lever-press task. (*E*) Outcome devaluation test. The number of lever presses during a 5-min test session under valued and devalued states was normalized to the number of lever presses in session 7. (*F*) Contingency degradation test. The number of lever presses in each 5-min block was normalized to the number of lever presses in session 12. (*G*) Session mean of the normalized number of lever presses (*n* = 8 for PBS, *n* = 9 for ITX). (*H*–*N*) Selective elimination of M1–DLS IT neurons and assessment of its behavioral effects. (*H*) Experimental strategy. NeuRet-IL-2Rα-GFP and NeuRet-RFP vectors were unilaterally injected into the Pn and the contralateral DLS, respectively, followed by an injection of either the LV-MCS or LV-FLEX-PE38 vector into M1 ipsilateral to the NeuRet-IL-2Rα-GFP injection. (*I*) Distribution of PT and IT neurons in M1. Sections through M1 were subjected to double immunohistochemistry with anti-GFP and anti-RFP antibodies. (*J*) Numbers of PT and IT neurons per section (*n* = 4 per group). (*K*–*N*) Behavioral assessment of *Tlx3-Cre* mice following M1–DLS IT neuron elimination. *Tlx3-Cre* mice received bilateral injections of the NeuRet-IL-2Rα-GFP vector into the Pn, followed by bilateral injections of the LV-MCS or LV-FLEX-PE38 injections into M1 and subjected to a series of lever-press tasks. (*K*) Acquisition of the lever-press task. (*L*) Outcome devaluation test. The number of lever presses was normalized as described for *E*. (*M*) Contingency degradation test. The number of lever presses in each block was normalized as described for *F*. (*N*) Session mean of the normalized number of lever presses (*n* = 9 per group). Data in *C*–*G* and *J*–*N* are presented as mean ± SEM; Statistical analyses were performed using unpaired *t*-test (*C*, *G, J* and *N*), two-way repeated-measures ANOVA followed by pairwise comparisons with Bonferroni correction (*D*, *F, K* and *M*), or paired *t*-test (*E* and *L*). See also *SI Appendix*, Table S1 for statistical details. \*\*\**P* < 0.001; ns, not significant. (Scale bars: *B* and *I*, 200 μm.)

For behavioral assessments, mice received bilateral injections of the NeuRet-IL-2Rα-GFP vector into the Pn, followed by ITX or PBS injections into M1. Both groups exhibited similar acquisition of the lever-press task (Fig. 4D).

In the outcome devaluation test, the normalized number of lever presses in the PBS-injected group did not differ significantly between the valued and devalued states, indicating insensitivity to outcome devaluation. In contrast, the ITX-injected group showed a significant reduction in lever pressing in the devalued state, indicating a failure to exhibit insensitivity to outcome devaluation (Fig. 4E). In the contingency degradation test, the normalized number of lever presses showed a similar pattern across blocks in both groups (Fig. 4F), and the session average did not differ significantly between the two groups (Fig. 4G). Thus, selective elimination of M1–DLS PT neurons impaired insensitivity to outcome devaluation without affecting performance in the contingency degradation test.

To eliminate IT neurons, we used *Tlx3-Cre* transgenic mice, which selectively express Cre recombinase in IT neurons (25, 26). The NeuRet-IL-2Rα-GFP and NeuRet-RFP vectors were unilaterally injected into the Pn and the contralateral DLS, respectively, followed by an injection of either a lentiviral vector encoding *Pseudomonas* exotoxin fragment PE38 (LV-FLEX-PE38) or a control lentiviral vector containing a multiple cloning site (MCS; LV-MCS) was injected into M1 ipsilateral to the NeuRet-IL-2Rα-GFP injection (Fig. 4H). Immunostaining of M1 sections revealed a reduction in the number of RFP-positive IT neurons in the LV-FLEX-PE38-injected group compared with the LV-MCS-injected group, whereas no apparent difference was observed in the number of GFP-positive PT neurons between the two groups (Fig. 4I). Quantitative analysis confirmed a significant reduction in the number of RFP-positive neurons in the LV-FLEX-PE38-injected group relative to the LV-MCS-injected group (Fig. 4J).

For behavioral assessment, *Tlx3-Cre* mice received bilateral injections of either the LV-FLEX-PE38 vector or the LV-MCS vector into M1. Both groups exhibited similar task acquisition (Fig. 4K). In the outcome devaluation test, the normalized number of lever presses did not differ significantly between the valued and devalued states in either group (Fig. 4L). In the contingency degradation test, the normalized number of lever presses also showed a similar pattern across blocks in the two groups (Fig. 4M), and the session average did not differ significantly between the groups (Fig. 4N). Selective elimination of M1–DLS IT neurons did not detectably alter behavioral insensitivity to either outcome devaluation or contingency degradation.

These findings suggest that PT and IT neuronal populations within the M1–DLS corticostriatal pathway have different functional roles in habitual behavior and that PT neurons contribute, at least in part, to the insensitivity to outcome devaluation associated with habit formation.

## Discussion

In the present study, we found that the M1–DLS corticostriatal pathway, but not S1–DLS pathway, plays an important role in habit formation, as assessed by the outcome devaluation and contingency degradation tests (Fig. 2 *K*–*R*). Our findings complement previous studies reporting that corticostriatal inputs from the PrL or OFC to the DMS contribute to the acquisition and expression of goal-directed actions (17–19). Together, these findings suggest functional specialization among corticostriatal pathways in goal-directed and habitual control during instrumental learning.

The motor cortex is implicated in planning, learning, and controlling motor behaviors (44, 45). Neuronal activity in M1, in particular, undergoes multiple plastic changes during instrumental learning processes (46, 47). Studies manipulating cortical neuronal activity under different experimental conditions have revealed diverse roles of the M1 in mice; for instance, optogenetic manipulation of M1 impairs the execution of learned movements in trained mice after initial training (48, 49), whereas the involvement of M1 declines after prolonged training (50, 51). Studies using chronic lesions have demonstrated that M1 is required for the acquisition or execution of various motor skills (51, 52). A recent study has also demonstrated that M1 neurons contribute to the execution of trained reaching behavior through modulation of striatal neuronal activity (53). In the present study, we identified a key role for M1 in habit formation, mediated, at least in part, by its corticostriatal projection to the DLS, extending previous findings on the involvement of the M1 in motor skill acquisition and execution (Fig. 2 *K*–*N*).

Within the M1, PT and IT neurons have distinct functional properties. A study using optogenetic identification of motor cortical neurons has shown that the firing activity of PT neurons is associated with contralateral movement, whereas IT neuronal activity is involved in bilateral movement during a head-restricted task in rats (27). A calcium-imaging-based brain-machine interface study reported that mice learn to control the activity of PT neurons faster than that of IT neurons during operant conditioning. This difference appears to be correlated with the location of the neurons in the cortex and local circuits (29). Although these studies have clarified the respective functional properties of PT and IT neurons, their potential involvement in habit formation remains unclear. In the present study, selective elimination of PT neurons in M1 resulted in a failure to exhibit the insensitivity to outcome devaluation observed in intact mice following instrumental training (Fig. 4E), whereas performance in the contingency degradation test was not significantly affected by selective elimination of either PT or IT neurons (Fig. 4 *F*, *G, M,* and *N*). These observations suggest that insensitivity to outcome devaluation and contingency degradation may depend, at least partly, on distinct neural mechanisms. In particular, our findings implicate PT neurons in the insensitivity to outcome devaluation associated with habit formation. PT neurons send collaterals to the DLS while also projecting broadly to subcortical structures, including the superior colliculus, pons, medulla, and spinal cord (21, 22). Whether and how the DLS-directed collaterals and other subcortical projections of PT neurons interact to contribute to habit formation remains an important question for future investigation. One possible explanation for why insensitivity to contingency degradation was not significantly affected by elimination of either PT or IT neurons is that the elimination was incomplete, allowing the remaining neurons to compensate and thereby preserve this insensitivity. Alternatively, PT and IT neurons may be functionally redundant in mediating the loss of sensitivity to contingency degradation. In this case, elimination of either population alone would leave this function intact, whereas a behavioral effect might emerge only after elimination of both populations.

Beyond the contributions of individual neuronal types, the role of the M1– DLS pathway in habit formation may be considered in relation to its potential interactions with other striatal input systems. Nigrostriatal dopaminergic signaling from the substantia nigra pars compacta (SNc) to the DLS has been implicated in habit formation during instrumental conditioning (54–56). Repeated dopaminergic activation also accelerates the emergence of habitual responding (57, 58). Corticostriatal glutamatergic signaling and nigrostriatal dopaminergic signaling interact to modulate the synaptic activity of striatal medium spiny neurons (59, 60). Glutamate receptor stimulation increases dopamine release from nigrostriatal terminals (61–63), and dopamine receptor stimulation alters striatal neuronal activity by modulating corticostriatal transmission (64, 65). Dopamine signaling also regulates the synaptic activity of striatal interneurons via presynaptic and postsynaptic mechanisms (66–68). The activity of these interneurons supports the regulation of synaptic plasticity within the striatal circuit (69, 70). In addition, the thalamostriatal system is essential for acquiring and executing motor skills through mechanisms involving synaptic plasticity (71). Together, these findings suggest that the M1– DLS corticostriatal glutamatergic system may interact with nigrostriatal dopaminergic and thalamostriatal systems to support habitual control. Whether and how the M1–DLS pathway interacts with either or both of these systems in habit formation remains to be addressed in the future.

Drug addiction has been proposed to involve alterations in the neural mechanisms underlying goal-directed and habitual drug seeking and is accompanied by substantial changes in multiple corticostriatal circuits (72, 73). An imbalance between goal-directed and habitual control systems, or altered habit formation, has been implicated in neuropsychiatric diseases, including obsessive-compulsive disorder and Tourette’s syndrome (74–77). The current findings may therefore provide a framework for understanding how alterations in corticostriatal pathways contribute to dysregulated habit formation in these conditions. Taken together, our findings identify the M1–DLS corticostriatal neurons as a key neural substrate for habit formation. This work establishes a foundational baseline for exploring how distinct cortical cell types and their subcortical networks interact to regulate habitual control in both health and disease.

### Materials and methods Experimental animals

Male C57BL/6J mice (CLEA Japan, Tokyo, Japan) and *Tlx3-Cre* mice (PL56 strain) (25), aged 8–12 weeks, were housed in groups of three or four per cage. The animals were maintained at 22 ± 2 °C and 60% humidity under a 12-h light/12-h dark cycle (lights on at 07:00), with free access to food and water. All animal care and experimental procedures were conducted in accordance with the institutional regulations for animal experiments at Fukushima Medical University. The study protocol was approved by the Institutional Animal Care and Use Committee of Fukushima Medical University.

### Viral vector preparation

NeuRet-IL-2Rα-GFP, NeuRet-Cre, and NeuRet-RFP vectors were prepared as described previously (78). The envelope plasmid contained cDNA encoding the fusion glycoprotein type E (FuG-E) under the control of the cytomegalovirus enhancer/chicken β-actin (CAG) promoter. FuG-E consists of the N-terminal region of the rabies virus glycoprotein extracellular domain and the membrane-proximal, transmembrane, and cytoplasmic domains of vesicular stomatitis virus glycoprotein (VSV-G). The transfer plasmids contained cDNAs encoding IL-2Rα-GFP, Cre, or RFP downstream of the murine stem cell virus (MSCV) promoter. LV-FLEX-PE38 and LV-MCS vectors were prepared as described previously (79), using an envelope plasmid expressing VSV-G cDNA under the control of the CAG promoter. Their transfer plasmid contained cDNA encoding PE38 or a multiple cloning site (MCS) downstream of the MSCV promoter. HEK293T cells were transfected with the transfer, envelope, and packaging plasmids using the calcium phosphate precipitation method. Viral particles were pelleted by centrifugation at 6,000 × *g* for 16–18 h at 4 °C and resuspended in PBS. For NeuRet vectors, the resuspended particles were applied to a Sepharose Q FF ion-exchange column (GE Healthcare, Chicago, IL) and eluted with a linear 0–1.5 M NaCl gradient. Peak fractions containing the viral particles were pooled and concentrated using a Vivaspin Turbo 4 (10,000 MWCO; Vivascience, Göttingen, Germany). For LV vectors, a 55% sucrose solution (w/v) in PBS, a 20% sucrose solution (w/v) in PBS, and the resuspended particles were sequentially layered. Centrifugation was performed at 100,000 × g for 2 h at 4 °C. The viral particles inside the visible white band were collected and concentrated using a Vivaspin Turbo 4 (10,000 MWCO; Vivascience). Viral genomic RNA was extracted using a NucleoSpin RNA Virus (Takara Bio Inc., Kusatsu, Japan), followed by reverse transcription and quantitative PCR with a Lenti-X qRT-PCR Titration Kit (Takara Bio Inc.). Quantitative PCR was performed on a StepOne Real-Time PCR System (Applied Biosystems, Tokyo, Japan) under the following conditions: 95 °C for 3 min (1 cycle), followed by 40 cycles of 95 °C for 15 s and 60 °C for 30 s.

AAV serotype 2 vectors were prepared using the AAV Helper-Free System (Agilent Technologies, Santa Clara, CA) as described previously (35). The transfer plasmids contained cDNAs encoding GFP, RFP, IL-2Rα-2A-ChR2-YFP, GluClα-mYFP or GluClβ-RFP, each flanked by FLEX cassette sequences, downstream of the CAG or elongation factor 1α (EF1α) promoter. HEK293T cells were transfected with the transfer and helper plasmids and a packaging plasmid carrying the AAV rep and cap genes using the calcium phosphate precipitation method. The crude viral lysate was purified via two rounds of CsCl gradient centrifugation, dialyzed, and concentrated with a Vivaspin Turbo 4 (10,000 MWCO; Vivascience). The viral genomic titers were determined by quantitative PCR as described previously (35).

### Intracranial surgery

Mice were anesthetized by intraperitoneal administration of a mixture of medetomidine (0.3 mg/kg), midazolam (4.0 mg/kg), and butorphanol (5.0 mg/kg) at a volume of 0.01 mL/g of body weight. Following anesthesia, the mice were secured in a stereotaxic frame (SR-5M-HT; Narishige, Tokyo, Japan). A midline scalp incision was made, and the periosteum was removed to expose the skull. The incisor bar of the stereotaxic frame was adjusted to position bregma and lambda at the same dorsoventral level. Small holes were drilled into the skull using an electric drill, and the dura was pierced with a stainless steel needle. A pulled glass micropipette was then inserted through these holes, and the tip was targeted to the specific brain regions.

The genomic titers of NeuRet-IL-2Rα-GFP, NeuRet-Cre, and NeuRet-RFP vectors were adjusted to 5.0 × 10^12^, 7.0 × 10^12^, and 5.0 × 10^12^ viral genomic copies/mL, to achieve comparable transgene expression levels in cortical areas. These vectors were injected into either the DLS (1.0 μL per site, 4 sites per hemisphere) or the Pn (1.0 μL per site, 1 site per hemisphere) through a glass micropipette connected to a microinfusion pump (ESP-32, Eicom, Kyoto, Japan) at a constant rate of 0.2 μL/min. Stereotaxic coordinates for the injection sites were determined using a mouse brain atlas (80), with the anteroposterior (AP) and mediolateral (ML) coordinates referenced to bregma and the dorsoventral (DV) coordinates measured from the dural surface. The AP/ML/DV coordinates (mm) were 1.0/2.0/3.1 (site 1), 1.0/2.0/2.4 (site 2), 0/2.5/3.1 (site 3), and 0/2.5/2.4 (site 4) for the DLS, and −4.0/0.5/5.3 for the Pn. The genomic titers for AAV-FLEX vectors ranged from 1.5 × 10^12^ to 4.5 × 10^12^ viral genomic copies/mL. These vectors were injected into M1 (0.5 μL per site, 4 sites per hemisphere) at a constant rate of 0.1 μL/min. The AP/ML/DV coordinates (mm) were 2.0/1.9/0.7 (site 1), 1.0/1.5/0.7 (site 2), 0/1.1/0.7 (site 3), and –0.8/1.0/0.7 (site 4). Genomic titers of LV-FLEX-PE38 or LV-MCS vectors were 4.0 × 10^11^ viral genomic copies/mL. These vectors were injected into M1 (1.0 μL per site, 7 sites per hemisphere) at a constant rate of 0.1 μL/min. The AP/ML/DV coordinates (mm) were 2.0/1.9/0.7 (site 1), 1.5/1.7/0.7 (site 2), 1.0/1.5/0.7 (site 3), 0.5/1.2/0.7 (site 4), 0/1.0/0.7 (site 5), –0.5/1.0/0.7 (site 6), and –1.0/1.1/0.7 (site 7).

The recombinant ITX, anti-Tac(Fv)-PE38 (81), was diluted to a final concentration of 40 μg/mL in PBS containing 0.1% mouse serum albumin. Four weeks after injection of the NeuRet-IL-2Rα-GFP vector, either ITX or PBS was injected into the M1 (7 sites per hemisphere) and S1 (5 sites per hemisphere), at 0.5 μL per site and a constant rate of 0.1 μL/min. The AP/ML/DV coordinates (mm) were 2.0/1.9/0.7 (site 1), 1.5/1.7/0.7 (site 2), 1.0/1.5/0.7 (site 3), 0.5/1.2/0.7 (site 4), 0/1.0/0.7 (site 5), –0.5/1.0/0.7 (site 6), and –1.0/1.1/0.7 (site 7) for M1, and 1.4/2.6/1.0 (site 1), 0.6/2.7/0.9 (site 2), –0.2/2.8/0.8 (site 3), – 1.0/2.5/0.7 (site 4), and –1.8/2.7/0.6 (site 5) for S1. Mice with mislocalized or insufficient cell ablation, low transgene expression, or extensive brain damage were excluded from the analysis.

### Histology and immunohistochemistry

Mice were anesthetized as described above and perfused transcardially with PBS, followed by 4% paraformaldehyde in 0.1 M phosphate buffer. The brains were dissected, post-fixed in the same fixative overnight, and cryoprotected by sequential immersion in 10%, 20%, and 30% sucrose in PBS solutions overnight at 4 °C . Serial coronal sections (30 μm thick) were cut consecutively using a cryostat (CM3050 S, Leica, Germany). Free-floating sections were blocked with 5% normal swine serum in PBS for 1 h at room temperature and incubated overnight at 4 °C with the following primary antibodies: rabbit anti-GFP (1:2000, FRONTIER INSTITUTE Co. Ltd, Hokkaido, Japan) and guinea pig anti-RFP (1:1000, FRONTIER INSTITUTE Co. Ltd). After rinsing in PBS, sections were incubated for 2 h at room temperature with the following species-specific secondary for either immunofluorescence (IF) or chromogenic (DAB) staining. For immunofluorescence, Alexa Fluor 488-conjugated anti-rabbit IgG (1:500, Thermo Fisher Scientific, Waltham, MA) and Cy3-conjugated anti-guinea pig IgG (1:500, Jackson ImmunoResearch Laboratories, West Grove, PA) were used. For DAB staining, biotinylated anti-rabbit IgG (1:500, Jackson ImmunoResearch Laboratories) was used, after which sections were rinsed again in PBS for 15 min and treated with an avidin-biotin-peroxidase complex kit (VECTASTAIN Elite ABC kit; Vector Laboratories Inc., Newark, CA).

Immunoreactivity was visualized by reacting the sections for 5–10 min with 0.05 M Tris-HCl buffer (pH 7.6) containing 0.05% 3,3′-diaminobenzidine tetrahydrochloride (DAB; DOJINDO, Kumamoto, Japan) and 0.3% H_2_O_2_ (FUJIFILM WAKO Pure Chemical, Osaka, Japan) at room temperature.

Sections were mounted onto gelatin-coated glass slides, dehydrated, and coverslipped. The IF- and DAB-stained sections were examined using a microscope (BZ-X810; Keyence, Osaka, Japan) capable of both fluorescence and brightfield imaging.

### *In vivo* electrophysiology and optogenetics

Mice were anesthetized as described above. Following anesthesia, the mice were secured in a stereotaxic frame (SR-5M; Narishige) using ear bars and a mouth-and-nose clamp (MA-6N; Narishige). After local anesthesia with 2% lidocaine, a midline scalp incision was made, and small holes were drilled through the skull. An optical fiber (FT200EMT; Thorlabs, Newton, NJ) was lowered into M1 at AP/ML/DV coordinates (mm) of 0.7–1.9/1.3–1.8/0.5–1.0, with placement varying across mice. A stainless-steel microelectrode (Unique Medical, Tokyo, Japan) was inserted into the DLS at AP/ML/DV coordinates (mm) of 0.5–1.0/2.3–3.0/1.5–2.3. Optical stimulation was delivered as 200 pulses of 465-nm light (11–13 mW, 50 ms per pulse, 1 Hz) using an optogenetic stimulation system (PlexBright, Plexon, Dallas, TX) controlled by an electronic stimulator (SEN-3301; Nihon Kohden, Tokyo, Japan). LFPs were recorded through the microelectrode and amplified using a bioelectric amplifier (MEG-5200; Nihon Kohden) via a miniature preamplifier (JH-110J; Nihon Kohden) and a signal selector (JB-120J; Nihon Kohden). The LFP signals were digitized at a sampling rate of 5 kHz with 16-bit resolution over an input range of –5 to 5 V and stored using a data acquisition system (PowerMicro1401-MKII and Spike2; Cambridge Electronic Design, Milton, UK). At the end of each experiment, electrolytic lesions were made along the electrode track by passing a positive current of 30 μA for 18 s through the microelectrode for subsequent histological verification. To mark the stimulation site, the optical fiber was removed, and the stainless-steel electrode was re-inserted along the optical fiber track to apply the lesioning current. The current ejected ferrocyanide ions from the electrode tip, allowing subsequent visualization of the lesions via the Prussian blue reaction.

### Slice electrophysiology

Mice were anesthetized as described above. Following anesthesia, the brains were dissected and coronal slices (300 μm thick) containing M1 were cut using a microslicer (PRO7, Dosaka, Kyoto, Japan) in ice-cold, oxygenated cutting Krebs’ solution containing (in mM) 120 choline chloride, 2.5 KCl, 26 NaHCO_3_, 1.25 NaH_2_PO_4_, 15 D-glucose, 1.3 ascorbic acid, 0.5 CaCl2, and 7 MgCl2. Slices were immediately transferred to a holding chamber containing standard Krebs’ solution composed of (mM) 124 NaCl, 3 KCl, 26 NaHCO_3_, 1 NaH_2_PO_4_, 2.4 CaCl2, 1.2 MgCl2, and 10 D-glucose (pH 7.4 when bubbled with 95% O_2_/5% CO_2_). Slices were then incubated in the holding chamber at room temperature (21–26 °C) for at least 1 h before recording. Neurons in M1 were visualized using an upright microscope (BX50WI, Olympus Optical Co., Tokyo, Japan) equipped with a 60× water-immersion objective. Fluorescently labeled corticostriatal neurons were identified by an appropriate fluorescence filter cube (U-MWIG3, Olympus Optical Co.). Patch pipettes were fabricated from standard-walled borosilicate glass capillaries (Harvard Apparatus, Cambridge, MA). For membrane potential recordings, the patch pipettes were filled with a K-gluconate-based internal solution containing (in mM) 120 K-gluconate, 6 NaCl, 5 CaCl2, 2 MgCl2, 0.2 K-EGTA, 10 K-HEPES, 2 Mg-ATP, and 0.3 Na-GTP (pH adjusted to 7.4 with 1 M KOH). Whole-cell patch-clamp recordings were obtained from the fluorescently labeled corticostriatal neurons using an amplifier (Axopatch 200B, Molecular Devices, San Jose, CA). Data were initially recorded on digital audiotapes using a DAT recorder (DC to 10 kHz; Sony, Tokyo, Japan) and subsequently digitized offline at 10 kHz after being low-pass filtered at 2 kHz with an 8-pole Bessel filter using pCLAMP 9 software (Molecular Devices).

### Behavioral analysis

Age-matched male littermate mice were used for the behavioral experiments. Mice were handled for 3 days before the beginning of behavioral testing, and food intake was restricted to maintain their body weight 80–90% of their *ad libitum* body weight during the operant conditioning tasks. Eight operant chambers (ENV-307W; Med Associates, St Albans, VT) were used for training. Each chamber was equipped with a pellet receptacle in the center of the front panel. Two retractable levers were mounted on either side of the receptacle, and a stimulus light was placed above each lever. Each chamber was enclosed in a sound-attenuating box and illuminated with a house light during the sessions. Equipment operation and behavioral data recording were controlled using Med-PC IV software (Med Associates).

Each mouse was acclimated to its assigned operant chamber and subsequently underwent a series of lever-press tasks in the same chamber throughout training. First, mice were trained to press a single lever (left or right) under an FR1 schedule, in which every lever press resulted in the delivery of one 20-mg food pellet (Bio-Serv, Frenchtown, NJ) as a reward. The FR1 session ended when mice earned 40 rewards (session 1). Mice were then trained under an RI30 schedule for 3 days (sessions 2–4) and an RI60 schedule for 3 days (sessions 5–7). Under the RI schedules, the first lever press after a randomly varying interval averaging 30 or 60 s resulted in the delivery of a food pellet. Each RI30 session ended after 30 min or delivery of 40 rewards, and each RI60 session ended after 30 min or delivery of 25 rewards, whichever occurred first in each case.

After acquiring lever pressing, mice underwent an outcome devaluation test (38, 39) on 2 consecutive days. On the first test day, mice had *ad libitum* access to laboratory chow (valued state) or the food pellets used as reinforcer (devalued state) for 1 h in individual consumption cages before a 5-min nonreinforced test session, during which the lever was extended but no reward was delivered (session 8). On the next day, the mice were exposed to the opposite prefeeding condition (valued or devalued) for 1 h and then underwent 5-min nonreinforced test session (session 9).

A 30-min contingency degradation (omission) test (40) was performed after mice had been retrained under the RI60 schedule for 3 consecutive days (sessions 10–12). During this test, a food pellet was delivered after every 20-s period without a lever press, and each lever press reset the timer. The test period was divided into six 5-min blocks for analysis. The number of lever presses during each test block was normalized to the number of lever presses recorded in session 12.

Spontaneous locomotor activity was measured with a movement analyzer equipped with photobeam sensors (SV-10; Tokyo Sangyou, Tokyo, Japan). The number of beam breaks was recorded over a 60-min period divided into six consecutive 10-min blocks. A rotarod test was also performed, in which mice were placed on an accelerating rotarod (ENV-575M; Med Associates), and the rotation speed was linearly increased from 4 to 40 rpm over a 5-min period in each trial. Mice underwent four trials per day with a 30-min intertrial interval for 2 consecutive days. The latency to fall from the rotating rod was measured to assess motor coordination and balance.

### Statistical analysis

Quantitative data are expressed as means ± SEM. Statistical comparisons were performed using IBM SPSS Statistics version 30 (IBM Corp., Armonk, NY). For multiple group comparisons, repeated-measures ANOVA followed by a *post hoc* Bonferroni multiple-comparisons test was used. For comparisons between two groups, unpaired or paired Student’s *t*-tests were applied, as detailed in *SI Appendix*, Table S1. Differences were considered statistically significant at *P* < 0.05.

### Data, materials, and software availability

All data supporting the findings of this study are included in the article and its *SI Appendix*, and can also be obtained from the corresponding author upon request without restrictions. Source data are available from Mendeley Data (https://data.mendeley.com/preview/4dsgy9gn96?a=cd83fc09-fbec-49be-a04c-2cd9eb13fda1).

## Supporting information

Supporting informations

## Acknowledgments

This work was supported by a Lotte Research Promotion Grant (No. LF000466 to S.K.) from Lotte Foundation; a Grant-in-Aid for Scientific Research (B) from the Japan Society for the Promotion of Science (No. 24K00509 to S.K., 25K02365 to K.K.); a grant from the Japan Agency for Medical Research and Development (No. 22dm0207113h0002 and 25wm0625103s0202 to K.K.); a Grant-in-Aid for Scientific Research on Transformative Research Areas (A) “Adaptive Circuit Census” (No. 21H05244 to K.K.) from the Ministry of Education, Culture, Sports, Science, and Technology of Japan; and a grant from the Core Research for Evolutional Science and Technology of the Japan Science and Technology Agency (to K.K.). We thank the following individuals and institutions for providing materials used in this study: Dr. I. Pastan (anti-Tac(Fv)-PE38); Dr. C. Gerfen (*Tlx3-Cre* mice); St. Jude Children’s Research Hospital (Dr. A. Nienhuis) and George Washington University (the HIV-1-based vector system); and Dr. B. Roth (plasmid DNA encoding hM4Di). We are grateful to M. Kikuchi, H. Hashimoto, and Y. Nakazato for technical support for vector preparation and to T. Kobayashi for the schematic illustrations.

## References

1. A. Dickinson, Actions and habits: the development of behavioural autonomy. Philos. Trans. R. Soc. Lond. B Biol. Sci. 308, 67–78 (1985).

2. B. W. Balleine, J. P. O’Doherty, Human and rodent homologies in action control: corticostriatal determinants of goal-directed and habitual action. Neuropsychopharmacology 35, 48–69 (2010).

3. C. D. Adams, A. Dickinson, Instrumental responding following reinforcement devaluation. Q. J. Exp. Psychol. 33, 109–121 (1981).

4. B. W. Balleine, A. Dickinson, Goal-directed instrumental action: contingency and incentive learning and their cortical substrates. Neuropharmacology 37, 407–419 (1998).

5. R. J. Dolan, P. Dayan, Goals and habits in the brain. Neuron 80, 312–325 (2013).

6. C. D. Adams, Variations in the sensitivity of instrumental responding to reinforcer devaluation. Q. J. Exp. Psychol. 34, 77–98 (1982).

7. M. E. Bouton, Habit and persistence. J. Exp. Anal. Behav. 121, 88–96 (2024).

8. H. H. Yin, B. J. Knowlton, The role of the basal ganglia in habit formation. Nat. Rev. Neurosci. 7, 464–476 (2006).

9. L. H. Corbit, Understanding the balance between goal-directed and habitual behavioral control. Curr. Opin. Behav. Sci. 20, 161–168 (2018).

10. B. W. Balleine BW, M. R. Delgado, O. Hikosaka, The role of the dorsal striatum in reward and decision-making. J. Neurosci. 27, 8161–8165 (2007).

11. J. Cox, I. B. Witten, Striatal circuits for reward learning and decision-making. Nat. Rev. Neurosci. 20, 482–494 (2019).

12. O. Hikosaka et al., Direct and indirect pathways for choosing objects and actions. Eur. J. Neurosci. 49, 637–645 (2019).

13. B. J. Hunnicutt et al., A comprehensive excitatory input map of the striatum reveals novel functional organization. Elife 5, e19103 (2016).

14. N. N. Foster et al., The mouse cortico–basal ganglia–thalamic network. Nature 598, 188–194 (2021).

15. H. H. Yin, B. J. Knowlton, B. W. Balleine, Lesions of dorsolateral striatum preserve outcome expectancy but disrupt habit formation in instrumental learning. Eur. J. Neurosci. 19, 181–189 (2004).

16. H. H. Yin, B. J. Knowlton, B. W. Balleine, Blockade of NMDA receptors in the dorsomedial striatum prevents action-outcome learning in instrumental conditioning. Eur. J. Neurosci. 22, 505–512 (2005).

17. C. M. Gremel, R. M. Costa, Orbitofrontal and striatal circuits dynamically encode the shift between goal-directed and habitual actions. Nat. Commun. 4, 2264 (2013).

18. C. M. Gremel, R. M. Costa, Premotor cortex is critical for goal-directed actions. Front. Comput. Neurosci. 7, 110 (2013).

19. G. Hart, L. A. Bradfield, B. W. Balleine, Prefrontal corticostriatal disconnection blocks the acquisition of goal-directed action. J. Neurosci. 38, 1311–1322 (2018).

20. J. K. O’Hare, et al., Pathway-specific striatal substrates for habitual behavior. Neuron 89, 472–479 (2016).

21. G. M. Shepherd, Corticostriatal connectivity and its role in disease. Nat. Rev. Neurosci. 14, 278–291 (2013).

22. Y. Kawaguchi, Pyramidal cell subtypes and their synaptic connections in layer 5 of rat frontal cortex. Cereb. Cortex 27, 5755–5771 (2017).

23. T. Kita, H. Kita, The subthalamic nucleus is one of multiple innervation sites for long-range corticofugal axons: a single-axon tracing study in the rat. J. Neurosci. 32, 5990–5999 (2012).

24. S. Jiang et al., Anatomically revealed morphological patterns of pyramidal neurons in layer 5 of the motor cortex. Sci. Rep. 10, 7916 (2020).

25. C. R. Gerfen, R. Paletzki, N. Heintz, GENSAT BAC cre-recombinase driver lines to study the functional organization of cerebral cortical and basal ganglia circuits. Neuron 80, 1368–1383 (2013).

26. B. M. Hooks et al., Topographic precision in sensory and motor corticostriatal projections varies across cell type and cortical area. Nat. Commun. 9, 3549 (2018).

27. S. Soma et al., Distinct laterality in forelimb-movement representations of rat primary and secondary motor cortical neurons with intratelencephalic and pyramidal tract projections. J. Neurosci. 37, 10904–10916 (2017).

28. A. F. Garcia et al., Distinct populations of cortical pyramidal neurons mediate drug reward and aversion. Nat. Commun. 12, 182 (2021).

29. N. Vendrell-Llopis et al., Diverse operant control of different motor cortex populations during learning. Curr. Biol. 32, 1616–1622.e5 (2022).

30. S. Kato et al., Neuron-specific gene transfer through retrograde transport of lentiviral vector pseudotyped with a novel type of fusion envelope glycoprotein. Hum. Gene Ther. 22, 1511–1523 (2011).

31. F. Schnütgen et al., A directional strategy for monitoring Cre-mediated recombination at the cellular level in the mouse. Nat. Biotechnol. 21, 562–565. (2012).

32. V. S. Sohal, F. Zhang, O. Yizhar, K. F. Deisseroth, Parvalbumin neurons and gamma rhythms enhance cortical circuit performance. Nature 459, 698–702. (2009).

33. A. Saunders, C. A. Johnson, B. L. Sabatini, Novel recombinant adeno-associated viruses for Cre activated and inactivated transgene expression in neurons. Front. Neural Circuits 6, 47. (2012).

34. S. Kato et al., Selective neural pathway targeting reveals key roles of thalamostriatal projection in the control of visual discrimination. J. Neurosci. 31, 17169–17179 (2011).

35. S. Kato et al., Action selection and flexible switching controlled by the intralaminar thalamic neurons. Cell Rep. 22, 2370–2382 (2018).

36. K. Kobayashi et al., Immunotoxin-mediated conditional disruption of specific neurons in transgenic mice. Proc. Natl. Acad. Sci. U.S.A. 92, 1132–1136 (1995).

37. K. Kobayashi, Controlled cell targeting system to study the brain neural circuitry. Neurosci. Res. 58, 118–123 (2007).

38. A. W. Johnson, A selective role for neuronal activity regulated pentraxin in the processing of sensory-specific incentive value. J. Neurosci. 27, 13430– 13435 (2007).

39. L. C. Hadjas, C. Lüscher, L. D. Simmler, Aberrant habit formation in the Sapap3-knockout mouse model of obsessive-compulsive disorder. Sci. Rep. 9, 12061 (2019).

40. A. L. Derusso et al., Instrumental uncertainty as a determinant of behavior under interval schedules of reinforcement. Front. Integr. Neurosci. 4, 17 (2010).

41. H. H. Yin et al., Dynamic reorganization of striatal circuits during the acquisition and consolidation of a skill. Nat. Neurosci. 12, 333–341 (2009).

42. P. F. Durieux, S. N. Schiffmann, A. de Kerchove d’Exaerde, Differential regulation of motor control and response to dopaminergic drugs by D1R and D2R neurons in distinct dorsal striatum subregions. EMBO J. 31, 640– 653 (2012).

43. W. Lerchner et al., Reversible silencing of neuronal excitability in behaving mice by a genetically targeted, ivermectin-gated Cl-channel. Neuron 54, 35–49 (2007).

44. J. N. Sanes, J. P. Donoghue, Plasticity and primary motor cortex. Annu. Rev. Neurosci. 23, 393–415 (2000).

45. J. Tanji, Sequential organization of multiple movements: involvement of cortical motor areas. Annu. Rev. Neurosci. 24, 631–651 (2001).

46. D. Huber et al., Multiple dynamic representations in the motor cortex during sensorimotor learning. Nature 484, 473–478 (2012).

47. A. J. Peters, S. X. Chen, T. Komiyama, Emergence of reproducible spatiotemporal activity during motor learning. Nature 510, 263–267 (2014).

48. J. Guo et al., Cortex commands the performance of skilled movement. Elife 4, e10774 (2015).

49. K. Morandell, D. Huber, The role of forelimb motor cortex areas in goal directed action in mice. Sci. Rep. 7, 15759 (2017).

50. E. J. Hwang et al., Disengagement of motor cortex from movement control during long-term learning. Sci. Adv. 5, eaay0001 (2019).

51. M. Heindorf, S. Arber, G. B. Keller, Mouse motor cortex coordinates the behavioral response to unpredicted sensory feedback. Neuron 99, 1040– 1054.e5 (2018).

52. R. Kawai et al., Motor cortex is required for learning but not for executing a motor skill. Neuron 86, 800–812 (2015).

53. M. A. Nicholas, E. A. Yttri, Motor cortex is responsible for motoric dynamics in striatum and the execution of both skilled and unskilled actions. Neuron 112, 3486–3501 (2024).

54. T. W. Robbins et al., Effects of dopamine depletion from the caudate-putamen and nucleus accumbens septi on the acquisition and performance of a conditional discrimination task. Behav. Brain Res. 38, 243–261 (1990).

55. A. Faure, U. Haberland, F. Condé, N. El Massioui, Lesion to the nigrostriatal dopamine system disrupts stimulus-response habit formation. J. Neurosci. 25, 2771–2780 (2005).

56. I. Márquez et al., Effects on goal directed behavior and habit in two animal models of Parkinson’s disease. Neurobiol. Learn Mem. 169, 107190 (2020).

57. A. Nelson, S. Killcross, Amphetamine exposure enhances habit formation. J. Neurosci. 26, 3805–3812 (2006).

58. A. J. D. Nelson, S. Killcross, Accelerated habit formation following amphetamine exposure is reversed by D1, but enhanced by D2, receptor antagonists. Front. Neurosci. 7, 76 (2013).

59. D. Centonze et al., Dopaminergic control of synaptic plasticity in the dorsal striatum. Eur. J. Neurosci. 13, 1071–1077 (2001).

60. D. J. Surmeier, J. Plotkin, W. Shen, Dopamine and synaptic plasticity in dorsal striatal circuits controlling action selection. Curr. Opin. Neurobiol. 19, 621−628 (2009).

61. D. W. Clow, K. Jhamandas, Characterization of L-glutamate action on the release of endogenous dopamine from the rat caudate-putamen. J. Pharmacol. Exp. Ther. 248, 722–728 (1989).

62. B. H. Westerink, M. Santiago, J. B. De Vries, The release of dopamine from nerve terminals and dendrites of nigrostriatal neurons induced by excitatory amino acids in the conscious rat. Naunyn. Schmiedebergs. Arch. Pharmacol. 345, 523–529 (1992).

63. A. Verma, B. Moghaddam, Regulation of striatal dopamine release by metabotropic glutamate receptors. Synapse 28, 220–226 (1998).

64. J. R. Brown, G. W. Arbuthnott, The electrophysiology of dopamine (D2) receptors: a study of the actions of dopamine on corticostriatal transmission. Neuroscience 10, 349–355 (1983).

65. M. Garcia-Munoz, S. J. Young, P. M. Groves, Terminal excitability of the corticostriatal pathway. I. Regulation by dopamine receptor stimulation. Brain Res. 551, 195–206 (1991).

66. T. Aosaki, K. Kiuchi, Y. Kawaguchi, Dopamine D1-like receptor activation excites rat striatal large aspiny neurons in vitro. J. Neurosci. 18, 5180–5190 (1998).

67. A Pisani et al., Activation of D2-like dopamine receptors reduces synaptic inputs to striatal cholinergic interneurons. J. Neurosci. 20, RC69 (2000).

68. E. Bracci, D. Centonze, G. Bernardi, P. Calabresi, Dopamine excites fast-spiking interneurons in the striatum. J. Neurophysiol. 87, 2190–2194 (2002).

69. M. Deffains, H. Bergman, Striatal cholinergic interneurons and cortico-striatal synaptic plasticity in health and disease. Mov. Disord. 30, 1014– 1025 (2015).

70. J. D. Berke, Functional properties of striatal fast-spiking interneurons. Front. Syst. Neurosci. 5, 45 (2011).

71. S. B. E. Wolff, R. Ko, B. P. Ölveczky, Distinct roles for motor cortical and thalamic inputs to striatum during motor skill learning and execution. Sci. Adv. 8, eabk0231 (2022).

72. B. J. Everitt, T. W. Robbins, Neural systems of reinforcement for drug addiction: from actions to habits to compulsion. Nat. Neurosci. 8, 1481– 1489 (2005).

73. B. J. Everitt, T. W. Robbins, Drug addiction: Updating actions to habits to compulsions ten years on. Annu. Rev. Psychol. 67, 23–50 (2016).

74. C. M. Gillan et al., Disruption in the balance between goal-directed behavior and habit learning in obsessive-compulsive disorder. Am. J. Psychiatry 168, 718–726 (2011).

75. C. M. Gillan et al., The role of habit in compulsivity. Eur. Neuropsychopharmacol. 26, 828–840 (2016).

76. C. Scholl, J. Baladron, J. Vitay, F. H. Hamker, Enhanced habit formation in Tourette patients explained by shortcut modulation in a hierarchical cortico-basal ganglia model. Brain Struct. Funct. 227, 1031–1050 (2022).

77. T. W. Weickert et al., Habit and skill learning in schizophrenia: evidence of normal striatal processing with abnormal cortical input. Learn Mem. 9, 430– 442 (2002).

78. S. Kato, K. Kobayashi, K. Kobayashi, Improved transduction efficiency of a lentiviral vector for neuron-specific retrograde gene transfer by optimizing the junction of fusion envelope glycoprotein. J. Neurosci. Methods. 227, 151–158 (2014).

79. S. Kato et al., Efficient gene transfer via retrograde transport in rodent and primate brains using a human immunodeficiency virus type 1-based vector pseudotyped with rabies virus glycoprotein. Hum. Gene Ther. 18, 1141−1151 (2007).

80. G. Paxinos, K. B. Franklin, The Mouse Brain in Stereotaxic Coordinates (Academic Press, 2019).

81. R. J. Kreitman et al., Recombinant immunotoxins containing anti-Tac(Fv) and derivatives of Pseudomonas exotoxin produce complete regression in mice of an interleukin-2 receptor-bearing human carcinoma. Blood 83, 426–434 (1994).

