## Supporting informations for "Habitual control of instrumental behavior driven by corticostriatal pathways originating from the primary motor cortex"

*To whom correspondence may be addressed.

**This file includes:**

Figures S1 to S6

Table S1

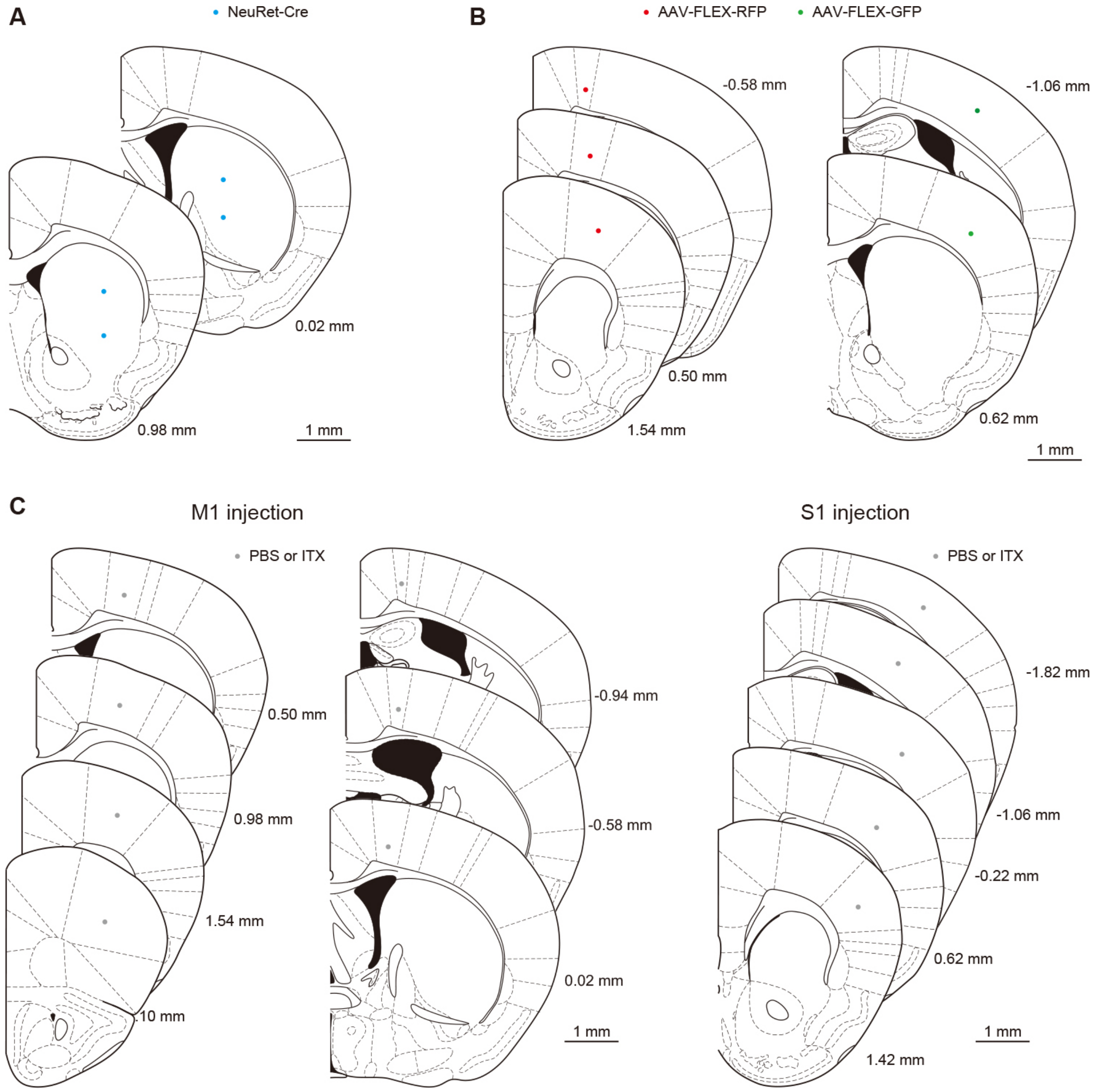


**Fig. S1. Sites of stereotaxic injections.** (*A*) Sites of NeuRet-Cre vector injections into the striatum. (*B*) Sites of AAV-FLEX-RFP and AAV-FLEX-GFP vector injections into the M1 and S1, respectively. (*C*) Sites of PBS or ITX injections into M1 and S1. Anteroposterior coordinates (mm) from bregma are indicated. Scale bars: *A*–*C*, 1 mm.

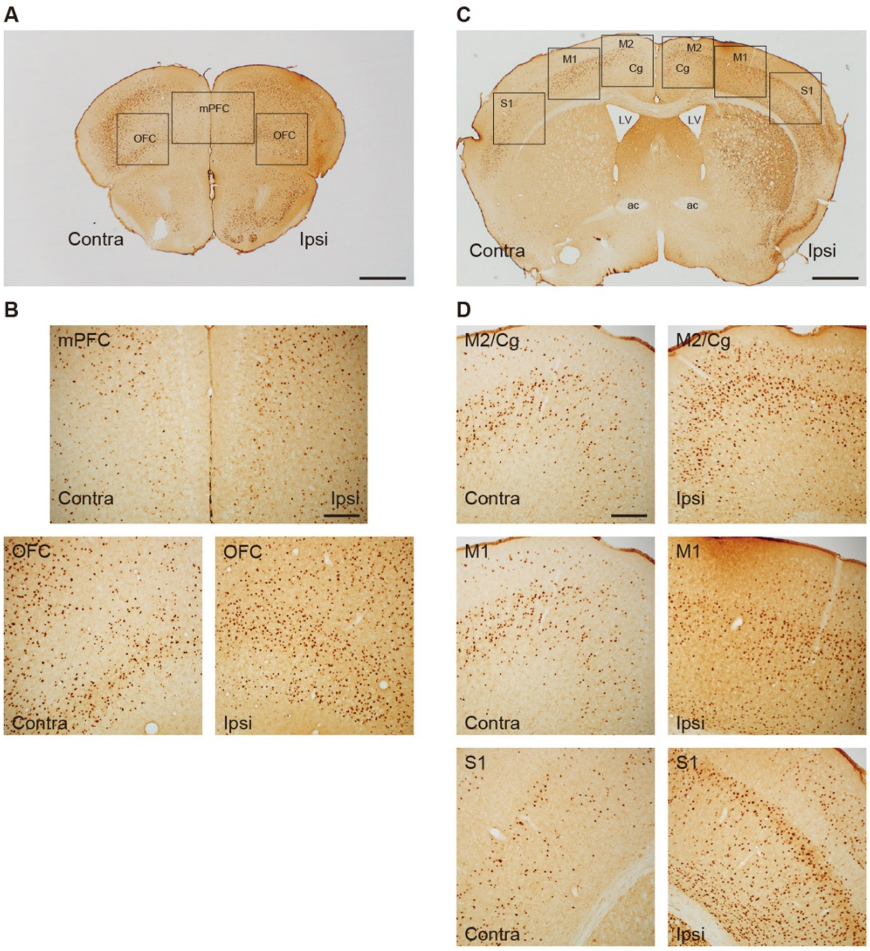


**Fig. S2. Cre immunohistochemistry following intrastriatal injection of the NeuRet-Cre vector.** Mice received a unilateral injection of the NeuRet-Cre vector into the dorsal striatum. Brain sections containing cortical areas were stained with an anti-Cre antibody. (*A* and *C*) Cre expression in the OFC and mPFC (*A*), and in the Cg, M2, M1, and S1 (*C*). (*B*) High-magnification images of the boxed regions indicated in (*A*). (*D*) High-magnification images of the square regions indicated in (*C*). Scale bars: *A* and *C,* 1 mm; *B* and *D*, 200 μm.

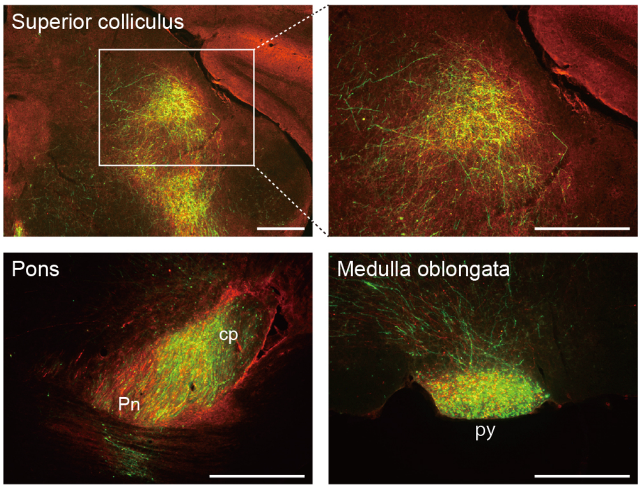


**Fig. S3. Axonal collaterals of corticostriatal neurons in subcortical regions contralateral to the vector injection side.** Mice received a unilateral injection of the NeuRet-Cre vector into the DLS, followed by ipsilateral injections of AAV-FLEX-RFP and AAV-FLEX-GFP vectors into M1 and S1, respectively. Brain sections were subjected to double immunohistochemistry with anti-RFP and anti-GFP antibodies. The upper panels show axonal collaterals of corticostriatal neurons in the contralateral superior colliculus. The upper right panel is a magnified view of the boxed region in the upper left panel. The lower panels show axonal collaterals in the contralateral pons and medulla oblongata. Pn, pontine nucleus; cp, cerebral peduncle; py, pyramidal tract. Scale bars: 400 μm.

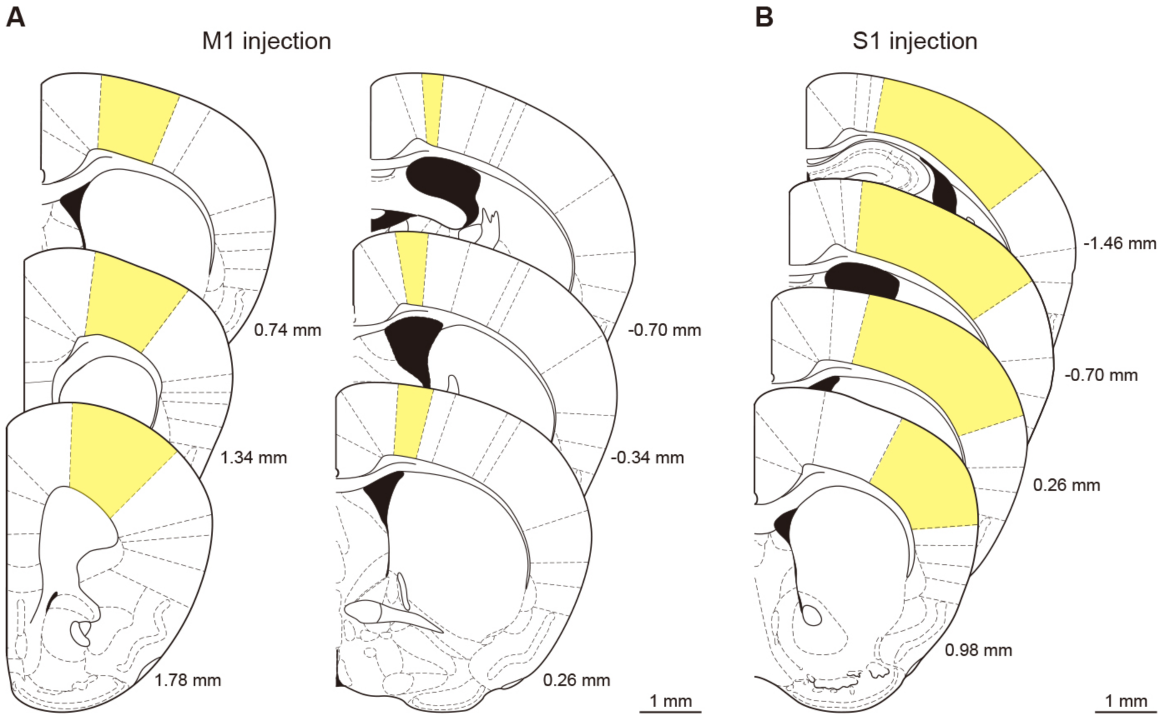


**Fig. S4. Schematic illustrations of brain sections used for cell counting.** The M1 (*A*) and S1 (*B*) regions are highlighted in yellow. Anteroposterior coordinates (mm) from bregma are indicated. Scale bars: *A* and *B*, 1 mm.

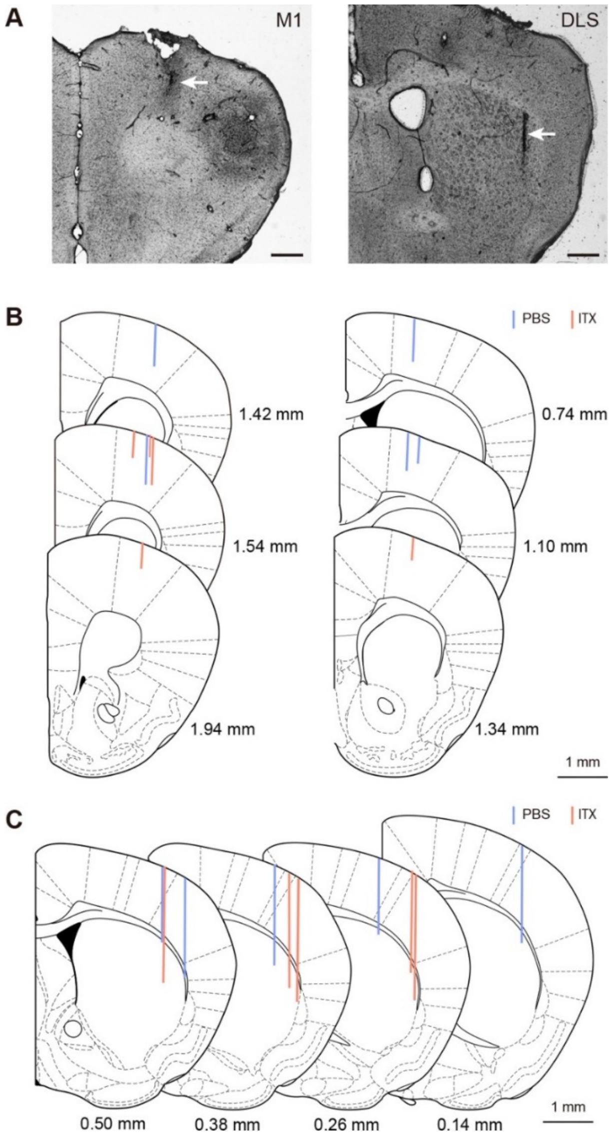


**Fig. S5. Placement sites of optical fibers and electrodes used for LFP recording.** After the recording experiments, an electrolytic lesion was made along the electrode track by passing a positive current (30 μA, 18 s). Brain sections (50 μm thick) containing M1 and DLS were prepared, and the marked sites were visualized using Prussian blue staining. (*A*) Representative sections showing the placement sites of an optical fiber in M1 (left) and an electrode in the DLS (right). The locations of their tips are indicated by arrows. (*B* and *C*) Schematic illustrations showing the placement sites of optical fibers in M1 (*B*) and electrodes in the DLS (*C*). Anteroposterior coordinates (mm) from bregma are indicated. Scale bars: *A*–*C*, 1 mm.

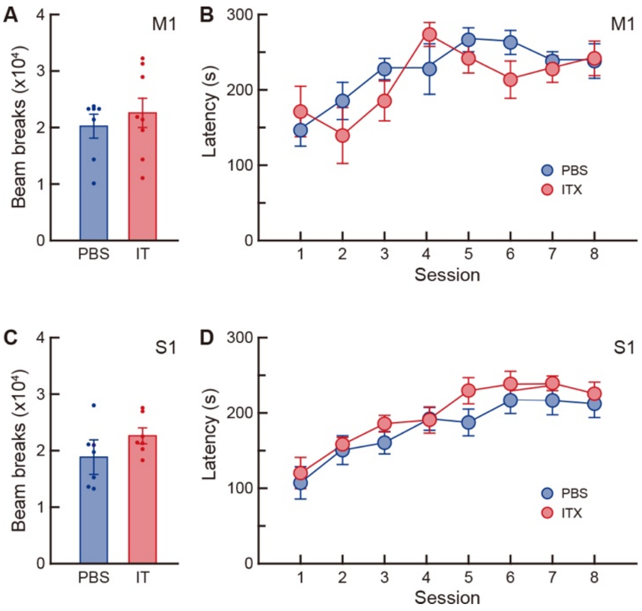


**Fig. S6. Motor performance after selective elimination of M1- or S1-derived corticostriatal neurons.** After selective elimination of M1- or S1-derived corticostriatal neurons using ITX, spontaneous locomotor activity and motor coordination were assessed, with PBS-injected mice serving as controls. (*A* and *B*) Spontaneous locomotor activity (*A*) and motor coordination (*B*) in mice with M1 injections. $n$ = 7 mice of PBS-group, $n$ = 8 mice of ITX-group. (*C* and *D*) Spontaneous locomotor activity (*C*) and motor coordination (*D*) in mice with S1 injections. $n$ = 7 mice each of PBS- and ITX-group for *C*, and $n$ = 8 mice each of PBS- and ITX-group for *D*. Data in *A*–*D* are presented as mean ± SEM; Statistical analyses were performed using unpaired $t$-test (*A* and *C*) or two-way repeated-measures ANOVA (*B* and *D*). See also *SI Appendix*, Table S1 for statistical details.
