## Supplementary material for "Habitual control of instrumental behavior driven by corticostriatal pathways originating from the primary motor cortex": TableS1

### Table S1. Summary of statistical analysis

| Figure | Experiment | Cohort | Outcome measure | n | p | Statistical test |
|---|---|---|---|---|---|---|
| Fig. 2B (M1) | ITX-mediated cell elimination | PBS/ITX in M1–DLS | Number of labeled cells in M1 | 6 and 6 mice | t(10) = 4.165<br>p < 0.01 (p = 0.00193) | unpaired *t*-test |
| Fig. 2B (S1) | ITX-mediated cell elimination | PBS/ITX in M1–DLS | Number of labeled cells in S1 | 6 and 6 mice | t(10) = 0.542,<br>p = 0.600 | unpaired *t*-test |
| Fig. 2E (M1) | ITX-mediated cell elimination | PBS/ITX in S1–DLS | Number of labeled cells in M1 | 6 and 6 mice | t(10) = 0.019<br>p = 0.985 | unpaired *t*-test |
| Fig. 2E (S1) | ITX-mediated cell elimination | PBS/ITX in S1–DLS | Number of labeled cells in S1 | 6 and 6 mice | t(10) = 12.009<br>p < 0.001 (p = 2.9007E-07) | unpaired *t*-test |
| Fig. 2J | LFP amplitude recording | PBS/ITX in M1–DLS | LFP amplitude after photostimulation | 5 and 5 mice | t(8) = 2.994<br>p < 0.05 (p = 0.01722) | unpaired *t*-test |
| Fig. 2K | Acquisition of lever-press task | PBS/ITX in M1–DLS | Number of lever presses during training | 7 and 8 mice | Group (PBS or ITX)<br>F(1,13) = 0.453<br>p = 0.513<br>Session<br>F(6,78) = 70.191<br>p < 0.001 (p = 2.14316E-29)<br>Group × Session<br>F(6,78) = 0.450<br>p = 0.843 | two-way ANOVA with repeated measures |
| Fig. 2L | Lever pressing during the outcome devaluation test | PBS/ITX and valued/devalued in M1–DLS | Number of lever presses normalized to session 7 | 7 and 8 mice | <PBS><br>t(6) = 0.247<br>p = 0.813<br><ITX><br>t(7) = 5.123<br>p < 0.01 (p = 0.00136) | paired *t*-test |
| Fig. 2M | Lever pressing during the contingency degradation test | PBS/ITX in M1–DLS | Number of lever presses normalized to session 12 | 7 and 8 mice | <Group>(PBS or ITX)<br>F(1,13) = 7.662<br>p < 0.05<br>(p = 0.01598)<br><Block><br>F(5,65) = 12.008<br>p < 0.001 (p = 3.00030E-08)<br><Group × Block><br>F(5,65) = 1.045<br>p = 0.399 | two-way ANOVA with repeated measures |
| Fig. 2N | Lever pressing during the contingency degradation test | PBS/ITX in M1–DLS | Number of lever presses normalized to session 12 | 7 and 8 mice | t(13) = 2.769<br>p < 0.05 (p = 0.01596) | unpaired *t*-test |
| Fig. 2O | Acquisition of lever-press task | PBS/ITX in S1–DLS | Number of lever presses during training | 10 and 11 mice | <Group>(PBS or ITX)<br>F(1,19) = 1.572<br>p = 0.225<br><Session><br>F(6,114) = 51.919<br>p < 0.001 (p = 2.30767E-30)<br><Group × Session><br>F(6,114) = 0.189<br>p = 0.979 | two-way ANOVA with repeated measures |
| Fig. 2P | Lever pressing during the outcome devaluation test | PBS/ITX and valued/devalued in S1–DLS | Number of lever presses normalized to session 7 | 10 and 11 mice | <PBS><br>t(9) = 0.562<br>p = 0.588<br><ITX><br>t(10) = 0.095<br>p = 0.926 | paired *t*-test |
| Fig. 2Q | Lever pressing during the contingency degradation test | PBS/ITX in S1–DLS | Number of lever presses normalized to session 12 | 10 and 11 mice | <Group>(PBS or ITX)<br>F(1,19) = 0.001<br>p = 0.980<br><Block><br>F(5,95) = 35.586<br>p < 0.001 (p = 2.32353E-20)<br><Group × Block><br>F(5,95) = 4.993<br>p < 0.001 (p = 0.00042)<br><br>*Post hoc* comparisons between PBS and ITX at each block<br>Block 1; t(19) = 1.989, p = 0.367<br>Block 2; t(19) = 0.606, p = 1.000<br>Block 3; t(19) = 1.624, p = 0.725<br>Block 4; t(19) = 1.109, p = 1.000<br>Block 5; t(19) = 0.289, p = 1.000<br>Block 6; t(19) = 1.519, p = 0.872 | two-way ANOVA with repeated measures |
| Fig. 2R | Lever pressing during the contingency degradation test | PBS/ITX in S1–DLS | Number of lever presses normalized to session 12 | 10 and 11 mice | t(19) = 0.659<br>p = 0.518 | unpaired *t*-test |
| Fig. 3C (Left) | Whole-cell patch-clamp recordings in brain slices | Before and after IVM application | Membrane potential | 5 mice | t(4) = 8.375<br>p < 0.01 (p = 0.00111) | paired *t*-test |
| Fig. 3C (Right) | Whole-cell patch-clamp recordings in brain slices | Before and after IVM application | Firing frequency | 5 mice | t(4) = 4.238<br>p < 0.05 (p = 0.01328) | paired *t*-test |

| | | | | | | |
|---|---|---|---|---|---|---|
| Fig. 3D | Acquisition of lever-press task | Vehicle/IVM in M1–DLS | Number of lever presses during training | 8 and 8 mice | <Group>(Vehicle or IVM)<br>F(1,14) = 0.321<br>p = 0.580<br><Session><br>F(6,84) = 36.226<br>p < 0.001 (p = 2.5012E-21)<br><Group × Session><br>F(6,84) = 0.871<br>p = 0.520 | two-way ANOVA with repeated measures |
| Fig. 3E | Lever pressing during the outcome devaluation test | Vehicle/IVM in M1–DLS | Number of lever presses normalized to session 7 | 8 and 8 mice | <Vehicle><br>t(7) = 0.071<br>p = 0.945<br><IVM><br>t(7) = 6.307<br>p < 0.001 (p = 4.0128E-4) | paired *t*-test |
| Fig. 3F | Lever pressing during the contingency degradation test | Vehicle/IVM in M1–DLS | Number of lever presses normalized to session 12 | 8 and 8 mice | <Group>(Vehicle or IVM)<br>F(1,14) = 4.701<br>p < 0.05 (p = 0.04786)<br><Block><br>F(5,70) = 18.083<br>p < 0.001 (p = 1.78131E-11)<br><Group × Block><br>F(5,70) = 0.596<br>p = 0.703 | two-way ANOVA with repeated measures |
| Fig. 3G | Lever pressing during the contingency degradation test | Vehicle/IVM in M1–DLS | Number of lever presses normalized to session 12 | 8 and 8 mice | t(14) = 2.741<br>p < 0.05 (p = 0.01591) | unpaired *t*-test |
| Fig. 4C (PT) | ITX-mediated cell elimination | PBS/ITX | Number of PT type neurons | 4 and 4 mice | t(6) = 15.101<br>p < 0.001 (p = 5.31783E-06) | unpaired *t*-test |
| Fig. 4C (IT) | ITX-mediated cell elimination | PBS/ITX | Number of IT type neurons | 4 and 4 mice | t(6) = 0.303<br>p = 0.772 | unpaired *t*-test |
| Fig. 4D | Acquisition of lever-press task | PBS/ITX in PT type neurons of M1–DLS | Number of lever presses during training | 8 and 9 mice | <Group>(PBS or ITX)<br>F(1,15) = 0.257<br>p = 0.619<br><Session><br>F(6,90) = 75.617<br>p < 0.001 (p = 5.37023E-33)<br><Group × Session><br>F(6,90) = 0.334<br>p = 0.917 | two-way ANOVA with repeated measures |
| Fig. 4E | Lever pressing during the outcome devaluation test | PBS/ITX and valued/devalued in PT type neurons of M1–DLS | Number of lever presses normalized to session 7 | 8 and 9 mice | <PBS><br>t(7) = 0.702<br>p = 0.505<br><ITX><br>t(8) = 7.650<br>p < 0.001 (p = 6.0188E-5) | paired *t*-test |
| Fig. 4F | Lever pressing during the contingency degradation test | PBS/ITX in PT type neurons of M1–DLS | Number of lever presses normalized to session 12 | 8 and 9 mice | <Group>(PBS or ITX)<br>F(1,15) = 0.223<br>p = 0.643<br><Block><br>F(5,75) = 20.183<br>p < 0.001 (p = 1.05974E-12)<br><Group × Block><br>F(5,75) = 0.279<br>p = 0.924 | two-way ANOVA with repeated measures |
| Fig. 4G | Lever pressing during the contingency degradation test | PBS/ITX in PT type neurons of M1–DLS | Number of lever presses normalized to session 12 | 8 and 9 mice | t(15) = 0.773<br>p = 0.451 | unpaired *t*-test |
| Fig. 4J (PT) | LV-FLEX-PE38-mediated cell elimination | LV-MCS/LV-FLEX-PE38 | Number of PT type neurons | 4 and 4 mice | t(6) = 0.073<br>p = 0.944 | unpaired *t*-test |
| Fig. 4J (IT) | LV-FLEX-PE38-mediated cell elimination | LV-MCS/LV-FLEX-PE38 | Number of IT type neurons | 4 and 4 mice | t(6) = 22.347<br>p < 0.001 (p = 5.25237E-07) | unpaired *t*-test |
| Fig. 4K | Acquisition of lever-press task | LV-MCS/LV-FLEX-PE38 in IT type neurons of M1–DLS | Number of lever presses during training | 9 and 9 mice | <Group>(LV-MCS or LV-FLEX-PE38)<br>F(1,16) = 0.130<br>p = 0.724<br><Session><br>F(6,96) = 40.698<br>p < 0.001 (p = 2.71417E-24)<br><Group × Session><br>F(6,96) = 1.664<br>p = 0.138 | two-way ANOVA with repeated measures |
| Fig. 4L | Lever pressing during the outcome devaluation test | LV-MCS/LV-FLEX-PE38 and valued/devalued in IT type neurons of M1–DLS | Number of lever presses normalized to session 7 | 9 and 9 mice | <LV-MCS><br>t(8) = 1.440<br>p = 0.188<br><LV-FLEX-PE38><br>t(8) = 2.090<br>p = 0.070 | paired *t*-test |

| | | | | | | |
|---|---|---|---|---|---|---|
| Fig. 4M | Lever pressing during the contingency degradation test | LV-MCS/LV-FLEX-PE38 in IT type neurons of M1–DLS | Number of lever presses normalized to session 12 | 9 and 9 mice | <Group>(LV-MCS or LV-FLEX-PE38)<br>F(1,16) = 3.359<br>p = 0.086<br><Block><br>F(5,80) = 20.539<br>p < 0.001 (p = 3.89913E-13)<br><Group × Block><br>F(5,80) = 2.131<br>p = 0.070 | two-way ANOVA with repeated measures |
| Fig. 4N | Lever pressing during the contingency degradation test | LV-MCS/LV-FLEX-PE38 in IT type neurons of M1–DLS | Number of lever presses normalized to session 12 | 9 and 9 mice | t(16) = 1.383<br>p = 0.186 | unpaired *t*-test |
| Fig. S6A | Spontaneous locomotor activity | PBS/ITX in M1–DLS | Number of beam breaks | 7 and 8 mice | t(13) = 0.714<br>p = 0.488 | unpaired *t*-test |
| Fig. S6B | Motor coordination and skill learning | PBS/ITX in M1–DLS | Latency to fall | 7 and 8 mice | <Group>(PBS or ITX)<br>F(1,13) = 0.340<br>p = 0.570<br><Session><br>F(7,91) = 8.451<br>p < 0.001 (p = 6.20313E-8)<br><Group × Session><br>F(7,91) = 1.716<br>p = 0.115 | two-way ANOVA with repeated measures |
| Fig. S6C | Spontaneous locomotor activity | PBS/ITX in S1–DLS | Number of beam breaks | 7 and 7 mice | t(12) = 1.591<br>p = 0.138 | unpaired *t*-test |
| Fig. S6D | Motor coordination and skill learning | PBS/ITX in S1–DLS | Latency to fall | 8 and 8 mice | <Group>(PBS or ITX)<br>F(1,14) = 1.360<br>p = 0.263<br><Session><br>F(7,98) = 17.248<br>p < 0.001 (p = 1.06763E-14)<br><Group × Session><br>F(7,98) = 0.457<br>p = 0.863 | two-way ANOVA with repeated measures |
